# Short-term dosage regimen for stimulation-induced long-lasting desynchronization

**DOI:** 10.1101/226134

**Authors:** Thanos Manos, Magteld Zeitler, Peter A. Tass

## Abstract

In this paper, we computationally generate hypotheses for dose-finding studies in the context of desynchronizing neuromodulation techniques. Abnormally strong neuronal synchronization is a hallmark of several brain disorders. Coordinated Reset (CR) stimulation is a spatio-temporally patterned stimulation technique that specifically aims at disrupting abnormal neuronal synchrony. In networks with spike-timing-dependent plasticity CR stimulation may ultimately cause an anti-kindling, i.e. an unlearning of abnormal synaptic connectivity and neuronal synchrony. This long-lasting desynchronization was theoretically predicted and verified in several pre-clinical and clinical studies. We have shown that CR stimulation with rapidly varying sequences (RVS) robustly induces an anti-kindling at low intensities e.g. if the CR stimulation frequency (i.e. stimulus pattern repetition rate) is in the range of the frequency of the neuronal oscillation. In contrast, CR stimulation with slowly varying sequences (SVS) turned out to induce an anti-kindling more strongly, but less robustly with respect to variations of the CR stimulation frequency. Motivated by clinical constraints and inspired by the spacing principle of learning theory, in this computational study we propose a short-term dosage regimen that enables a robust anti-kindling effect of both RVS and SVS CR stimulation, also for those parameter values where RVS and SVS CR stimulation previously turned out to be ineffective. Intriguingly, for the vast majority of parameter values tested, spaced multishot CR stimulation with demand-controlled variation of stimulation frequency and intensity caused a robust and pronounced anti-kindling. In contrast, spaced CR stimulation with fixed stimulation parameters as well as singleshot CR stimulation of equal integral duration failed to improve the stimulation outcome. In the model network under consideration, our short-term dosage regimen enables to robustly induce long-term desynchronization at comparably short stimulation duration and low integral stimulation duration. Currently, clinical proof of concept is available for deep brain CR stimulation for Parkinson’s therapy and acoustic CR stimulation for tinnitus therapy. Promising first in human data is available for vibrotactile CR stimulation for Parkinson’s treatment. For the clinical development of these treatments it is mandatory to perform dose-finding studies to reveal optimal stimulation parameters and dosage regimens. Our findings can straightforwardly be tested in human dose-finding studies.

## Introduction

To establish a pharmacological treatment for clinical use, in humans typically a 4-phase sequence of clinical trials is performed (Friedman et al., 2010). In pre-clinical studies pharmacokinetic, toxicity and efficacy are studied in non-human subjects. In first in human-studies (phase I) safety and tolerability of a drug are studied in healthy volunteers. Proof of concept studies (phase IIA) determine whether a drug can have any efficacy, whereas dose-finding studies (phase IIB) are performed to reveal optimum dose at which a drug has biological activity with minimal side-effects. Effectiveness and the clinical value of a new intervention are studied in a randomized controlled trial (phase III), compared with state of the art treatment, if available. Finally, post-marketing surveillance trials (phase IV) are performed to detect rare or long-term adverse effects within a much larger patient population and over longer time periods. There might also be combinations of different phases.

In principle, this 4-phase pattern is also valid for medical technology, e.g. neuromodulation technologies. However, if neuromodulation technologies aim at the control of complex dynamics of e.g. neural networks, different parameters and dosage regimens may have complex, non-linear and even counterintuitive effects, see e.g. (Gao et al., 2014;Popovych et al., 2015;Gates and Rocha, 2016;Zañudo et al., 2017;Zhang et al., 2017). This computational paper illustrates how computational modelling can be used to generate hypotheses for dose-finding studies. In general, performing dose-finding studies simply by trial and error may be impossible because of the substantial parameter space to be tested, with trial durations and related costs getting out of hands.

The development of proper dosage strategies and regimens enables favorable compromises between therapeutic efficacy and detrimental factors such as side-effects or treatment duration. This is relevant, e.g. for the development of pharmaceutical (Williams, 1992;Bertau et al., 2008;Peters, 2012;Dash et al., 2014) or radiation therapy (Symonds et al., 2012). Deep brain stimulation (DBS) is the standard treatment of medically refractory movement disorders (Benabid et al., 1991;Krack et al., 2003;Deuschl et al., 2006). The clinical (Temperli et al., 2003) and electrophysiological (Kühn et al., 2008;Bronte-Stewart et al., 2009) effects of standard high-frequency (HF) DBS occur only during stimulation and cease after stimulation offset.

Coordinated reset (CR) stimulation (Tass, 2003a;Tass, 2003b) was computationally developed to specifically counteract abnormal neuronal synchrony by desynchronization. CR stimulation uses sequences of stimuli delivered to neuronal sub-populations engaged in abnormal neuronal synchronization (Tass, 2003a;Tass, 2003b). As shown computationally, in neuronal populations with spike-timing-dependent plasticity (STDP) (Gerstner et al., 1996;Markram et al., 1997;Bi and Poo, 1998) CR stimulation may have long-lasting, sustained effects (Tass and Majtanik, 2006;Hauptmann and Tass, 2007;Popovych and Tass, 2012). This is because in the presence of STDP, CR stimulation reduces the rate of coincidences. Accordingly, the network may be shifted from an attractor with abnormal synaptic connectivity and abnormal neuronal synchrony to an attractor with weak connectivity and synchrony (Tass and Majtanik, 2006;Hauptmann and Tass, 2007;Popovych and Tass, 2012). This process was termed anti-kindling (Tass and Majtanik, 2006).

Abnormal neuronal synchronization has been shown to be associated with a number of brain diseases, for example, Parkinson’s disease (PD) (Lenz et al., 1994;Nini et al., 1995;Hammond et al., 2007), tinnitus (Ochi and Eggermont, 1997;Llinas et al., 1999;Weisz et al., 2005;Eggermont and Tass, 2015), migraine (Angelini et al., 2004;Bjørk and Sand, 2008). In parkinsonian nonhuman primates it was shown that electrical CR stimulation of the subthalamic nucleus (STN) has sustained, long-lasting after-effects on motor function (Tass et al., 2012b;Wang et al., 2016). In contrast, long-lasting after-effects were not observed with standard HF DBS (Tass et al., 2012b;Wang et al., 2016). For instance, unilateral CR stimulation of the STN of parkinsonian MPTP monkeys, delivered for only 2 h per day during 5 consecutive days led to significant and sustained bilateral therapeutic after-effects for at least 30 days, whereas standard HF DBS had no after-effects (Tass et al., 2012b). By the same token, cumulative and lasting after-effects of electrical CR stimulation of the STN were also observed in PD patients (Adamchic et al., 2014).

HF DBS may not only cause side effects by electrical current spreading outside of the target region, but also by chronic stimulation of the target itself or by functional disconnection of the stimulated structure (Ferraye et al., 2008;Moreau et al., 2008;van Nuenen et al., 2008). Accordingly, it is key to reduce the integral stimulation current. Electrical CR stimulation of the STN employs significantly less current compared to HF DBS (Tass et al., 2012b;Adamchic et al., 2014;Wang et al., 2016). However, to further improve the CR approach, in a previous computational study the spacing principle (Ebbinghaus et al., 1913) was used to achieve an anti-kindling at *subcritical intensities*, i.e. particularly weak intensities rendering permanently delivered CR stimulation ineffective (Popovych et al., 2015). According to the spacing principle (Ebbinghaus et al., 1913), learning effects can be improved by repeated stimuli spaced by pauses as opposed to delivering a massed stimulus in a single long stimulation session. The spacing principle was investigated on different levels, ranging from behavioral and cognitive (Cepeda et al., 2006;Pavlik and Anderson, 2008;Cepeda et al., 2009;Xue et al., 2011;Kelley and Whatson, 2013) to neuroscientific and molecular (Itoh et al., 1995;Frey and Morris, 1997;Menzel et al., 2001;Scharf et al., 2002;Naqib et al., 2012). Computationally it was demonstrated that the spacing principle can also be applied to unlearn unwanted, upregulated synaptic connectivity at subcritical stimulation intensities (Popovych et al., 2015). In principle, the results were intriguing, but required rather long pauses and total stimulation durations (Popovych et al., 2015). Spaced CR stimulation at subcritical intensities might possibly be applied to CR DBS. However, for clinical applications, in particular, for non-invasive applications of CR stimulation (Popovych and Tass, 2012), such as acoustic CR stimulation for tinnitus (Tass et al., 2012a) or vibrotactile stimulation for PD (Syrkin-Nikolau et al., 2017;Tass, 2017), it is crucial to achieve therapeutic effects within a reasonable amount of time. Applications of non-invasive medtech devices typically rely on the patients’ compliance and should favorably require short stimulation durations. Accordingly, we here set out to apply the spacing principle to CR stimulation at *supercritical intensities*, i.e. intensities that enable an anti-kindling for moderate stimulation duration and properly selected stimulation frequencies. The overall goal of this study is to design short-term dosage regimen that improve CR stimulation efficacy, while keeping the integral amount of stimulation as well as the overall duration of the protocols at comparably low levels.

In (Manos et al., 2017) we studied the influence of the CR stimulation frequency and the intensity on the outcome of CR stimulation with *Rapidly Varying Sequences* (RVS) and *Slowly Varying Sequences* SVS (Zeitler and Tass, 2015). CR stimulation consists of sequences of stimuli delivered to each sub-population (Tass, 2003a;Tass, 2003b). For RVS CR stimulation, the CR sequence is randomly varied from one CR stimulation period to another (Tass and Majtanik, 2006). Conversely, SVS CR stimulation is characterized by repeating a sequence for a number of times before randomly switching to the next sequence (Zeitler and Tass, 2015). In (Manos et al., 2017) we demonstrated that the efficacy of singleshot CR stimulation with moderate stimulation duration depends on the stimulation parameters, in particular, on the intensity as well as the relationship between CR stimulation frequency and intrinsic firing rates. RVS CR stimulation turned out to induce pronounced long-lasting desynchronization, e.g. at weak intensities and CR stimulation frequencies in a certain range around the neurons’ intrinsic firing frequencies. In contrast, SVS CR stimulation enabled even more pronounced anti-kindling, however, at the cost of a significantly stronger dependence of the stimulation outcome on the CR stimulation frequency.

Dosage regimen design is an integral part of pharmacokinetic methodology, aiming at an optimization of drug delivery and effects (Williams, 1992). By a similar token, we hypothesize that appropriate dosage regimens might further enhance the efficacy of RVS and SVS CR stimulation. To probe different dosage regimens, we here consider different stimulation singleshot and multishot CR stimulation protocols. *Protocols A* and *B* have identical integral stimulation duration, whereas *Protocols C* and *D* may require less stimulation.

### Protocol A: Spaced multishot CR stimulation with fixed stimulation parameters

Instead of one singleshot CR stimulation we deliver the identical CR shot five times, where the duration of each single pause equals the duration of each identical singleshot. Intersecting singleshot stimuli by pauses to increase stimulation efficacy, resembles the so-called *spacing principle*, a learning-related mechanism that is well-established in psychology (Ebbinghaus et al., 1913), education (Kelley and Whatson, 2013), and neuroscience (Naqib et al., 2012). According to the spacing principle, learning effects can be enhanced by delivering a stimulus in a spaced manner, as opposed to administering a massed stimulus in a single long stimulation session. Computationally, it was shown that subcritical CR stimulation at subcritical (ineffective) intensities may become effective if intersected by rather long pauses and delivered sufficiently often, e.g. eight times (Popovych et al., 2015). However, shorter pauses were not sufficient (Popovych et al., 2015). As yet, spaced CR stimulation at supercritical intensities was not studied. Here, we focus on comparably short stimulation protocols. Accordingly, we use CR stimulation of sufficient intensity and deliver five single CR shots intersected by pauses. We consider a symmetric dosage regimen, with identical duration of single shots and pauses.

### Protocol B: Long singleshot CR stimulation with fixed stimulation parameters

To assess the impact of the spacing principle, as a control condition we simply stimulate five times longer, without any pause and with stimulation parameters kept constant. *Protocol B* is shorter, but employs the same integral stimulation duration as *Protocol A*.

### Protocol C: Spaced multishot CR stimulation with demand-controlled variation of the CR stimulation frequency and intensity

As in *Protocol A*, we deliver spaced CR stimulation comprising five identical CR shots, intersected by pauses, where all shots and pauses are of equal duration. However, at the end of each CR shot we monitor the stimulation effect and perform a three-stage control: (i) If no pronounced desynchronization is achieved, the CR stimulation frequency of the subsequent CR shot is mildly varied by no more than ±3%. (ii) If an intermediate desynchronization is observed, the CR stimulation frequency remains unchanged and CR stimulation is continued during the subsequent shot. (iii) If a pronounced desynchronization is achieved, no CR stimulation is delivered during the subsequent shot. Note, for stage (i) we do not adapt the CR stimulation frequency to a measured quantity. We consider two different variation types employed for stage (i): with regular and with random variation of the CR stimulation frequency. Regular variation means to increase or decrease the CR stimulation frequency in little unit steps. In contrast, random variation stands for randomly picking the CR stimulation frequency from a restricted interval.

### Protocol D: Long singleshot CR stimulation with demand-controlled variation of the stimulation frequency

To assess the specific pausing-related impact of the evolutionary spacing principle, as a direct control condition we perform *Protocol C* without pauses. To this end, we string five CR shots together, without pauses, and evaluate the stimulation effect at the end of each CR shot. If no pronounced desynchronization is achieved, the CR stimulation frequency is slightly varied by no more than ±3% for the subsequent CR shot. During each single CR shot stimulation parameters are kept constant. Only from one CR shot to the next the CR stimulation frequency can be varied. Overall, *Protocol D* is shorter than *Protocol C*, but uses the same integral stimulation duration as in *Protocols A-C*.

CR stimulation and, especially, SVS CR stimulation has pronounced periodic characteristics. Accordingly, the CR stimulation frequency turned out to be a sensitive parameter, in particular, for SVS CR stimulation [see (Manos et al., 2017)]. For this reason, for stage (i) of *Protocol C* and *D* we perform a demand-controlled variation of the CR stimulation frequency to prevent from, e.g. unfavorable resonances or phase locking dynamics. Note these demand-controlled changes of the CR stimulation frequency are mild and hardly change the networks’ firing rates.

In this study, we test the performance of the different *Protocols A-D* by selecting unfavorabl stimulation parameters, which render CR stimulation ineffective according to (Manos et al., 2017) By design, *Protocols C* and *D* work well for all parameter pairs (*K, T_s_*) related to effective singleshot CR stimulation. In that case, CR stimulation actually ceases due to lack of demand. Note, in all four stimulation protocols we keep the stimulation intensity fixed. Only *Protocols C* and *D* require feedback of the stimulation outcome

This paper is organized as follows: in the *Materials and Methods* section we briefly describe the computational model, the neural network (and its initialization), the synaptic plasticity rule, the CR stimulation, the analysis methods used throughout the paper as well as the summary of the CR frequency and intensity global trends which were thoroughly studied in (Manos et al., 2017). In the *Results* section, we present all the different *Protocols* in detail and our main findings regarding their comparison. Finally, in the *Discussion* section, we discuss our findings and set this work in more general perspective, related to medical applications.

## Materials and Methods

### Model and Network Description

In this study we use the conductance-based Hodgkin-Huxley neuron model (Hodgkin and Huxley, 1952) for the description of an ensemble of spiking neurons. The set of equations and parameters read [see (Hansel et al., 1993;Popovych and Tass, 2010)]:

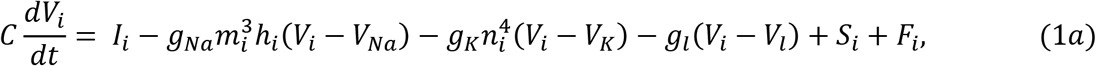

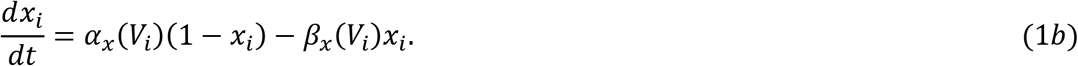

Variable *V_i_* denotes the membrane potential of neuron *i* (*i* = 1,…, *N*), while the variable *x* stands for the three gating variables *m, n* and *h*. The *α_x_* and *β_x_* variables are described in the standard model definition [see (Manos et al., 2017)]. The network consists of *N* = 200 neurons placed on a ring. The constant sodium, potassium and leak reversal potentials and the maximum conductance per unit area are (*V_Na_, g_Na_*) = (50 mV, 120 mS/cm^2^), (*V_K_,g_K_*) = (−77mV, 36 mS/cm^2^) and (*V_l_,g_l_*) = (−54.4 mV, 0.3 mS/cm^2^), while the constant membrane capacitance is *C* = 1 μF/cm. *I_i_* denotes the constant depolarizing current injected into neuron *i*, regulating the intrinsic firing rate of the uncoupled neurons. For the realization of different initial networks, we used the same random initial conditions drawn from uniform distributions as used in (Manos et al., 2017), i.e. *I_i_* ∈ [*I*_0_ – *σ_I_, I*_0_ + *σ_l·_*] (*I*_0_ = 11.0 μA/cm^2^ and *σ_l_* = 0.45 μA/cm^2^), *h_j_,m_i_,n_i_* ∈ [0,1] and *V_i_* ∈ [−65,5] mV. In addition, in order to model variations of the model parameters (see *Discussion* and *Supplementary Material*), we add a sinusoidal external current input of the form *I_var_* = *A* · sin(2*π* · *f* · *t*) to the right-hand side of Equation 1a, where *f* and *A* are the frequency and the amplitude of the signal respectively.

The initial values of the neural synaptic weights *c_ij_* are picked from a normal distribution *N*(μ_*c*_ = 0.5 mS/cm^2^, *σ_c_* = 0.01 mS/cm^2^) as in (Manos et al., 2017) [see (Popovych and Tass, 2012;Zeitler and Tass, 2015) for details]. *S_i_*(*t*) denotes the internal synaptic input within the network to neuron *i*. The neurons interact via excitatory and inhibitory chemical synapses *s_i_*, by means of the postsynaptic potential (PSP) *s_i_* which is triggered by a spike of neuron *i* (Gerstner et al., 1996;Izhikevich, 2010) and modelled using an additional equation [see (Golomb and Rinzel, 1993;Terman et al., 2002)]:

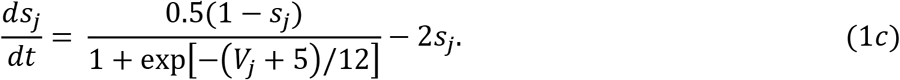

Initially we draw *s_i_* ∈ [0,1] (randomly from a uniform distribution). The coupling term *S_i_* from Equation 1a [see (Popovych and Tass, 2012)] contains a weighted ensemble average of all postsynaptic currents received by neuron *i* from the other neurons in the network: 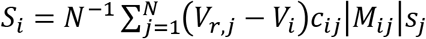 where *V_r,j_* is the reversal potential of the synaptic coupling (20 mV for excitatory and −40 mV for inhibitory coupling), and *c_ij_* is the synaptic coupling strength from neuron *j* to neuron *i*. There are no neuronal self-connections within the network (*c_ii_* = 0 mS/cm^2^).

The variable:

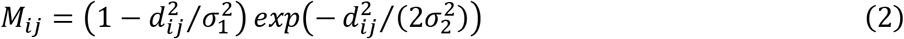

describes the spatial profile of coupling between neurons *i* and *j* and is of a Mexican hat-type (Wilson and Cowan, 1973;Dominguez et al., 2006;de la Rocha et al., 2008) with strong short-range excitatory (*M_ij_* > 0) and weak long-range inhibitory interactions (*M_ij_* < 0). Here *d_ij_* = *d*|*i* – *j*| is the distance between neurons *i* and *j*, while *d* = *d*_0_/(*N* – 1) determines the distance on the lattice between two neighboring neurons within the ensemble. *d*_0_ is the length of the neuronal chain (*d*_0_ = 10). *σ*_1_ = 3.5, and *σ*_2_ = 2.0. In order to limit boundary effects, we consider that the neurons are distributed in such a way that the distance *d_ij_* is taken as: *d* · *min*(*i* – *j*|,*N* – |*i* – *j*|) for *i,j* > *N*/2.

### Spike-Timing-Dependent Plasticity

The synaptic weights *c_ij_* are dynamical variables that depend on the time difference, Δ*t_ij_* = *t_i_* – *t_j_*, between the onset of the spikes of the post-synaptic neuron *i* and the pre-synaptic neuron *j*, denoted by *t_i_* and *t_j_*, according to (Bi and Poo, 1998;Popovych and Tass, 2012):

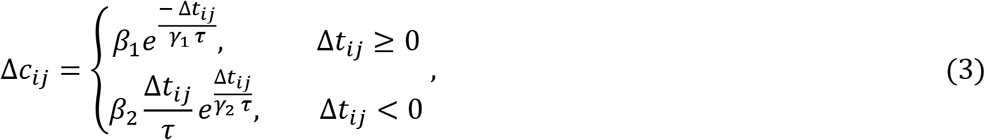

with parameters *β*_1_ = 1, *β*_2_ = 16, *γ*_1_ = 0.12, *γ*_2_ = 0.15, *τ* = 14 ms and *δ* = 0.002, while the values of *c_ij_* are confinded to the interval [0,1] mS/cm^2^ for both excitatory and inhibitory synapses and, hence, remain bounded.

### Coordinated Reset Stimulation

The term *F_i_* in Equation 1a represents the current induced in neuron *i* by the CR stimulation delivered at *N_s_* = 4 stimulations sites, equidistantly placed at the positions of neurons *i* = 25,75,125,175 (Tass, 2003b). One stimulation site was active during *T_s_*/*W_s_*, while the other stimulation sites were inactive during that time window. After this, another stimulation site was active during the next *T_s_*/*N_s_* window. All stimulation sites were stimulated exactly once within one CR stimulation period of duration *T_s_*. The spatiotemporal activation pattern of stimulation sites is represented by the indicator functions *ρ_k_*(*t*) (*kϵ* {1, …, *N*}), taking the value 1 when the *k^th^* stimulation site is active at t and 0 else. The stimulation signals induced single brief excitatory post-synaptic currents. The evoked time-dependent normalized conductances of the postsynaptic membranes are represented by *α*-functions given in (Popovych and Tass, 2010) as 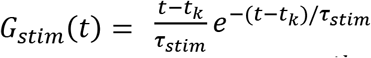, *t_k_* ≤ *t* ≤ *t*_*k*+_. *τ_stim_* = *T_s_*/(6*N_s_*) denotes the time-to-peak of *G_stim_*, and *t_k_* is the onset of the *k^th^* activation of the stimulation site. Note, *τ_stim_* determines the onset timing of each single signal as well as its duration. The spatial spread of the induced excitatory postsynaptic currents in the network is defined by the quadratic spatial decay profile 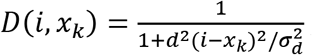, a function of the difference between the index of neuron *i* and the index *x_k_* of the neuron at stimulation site *k. d* is the lattice distance between two neighboring neurons, and *σ_d_* = 0.08 the spatial decay rate of the stimulation current [see (Popovych and Tass, 2010) for details]. Thus, the total stimulation current from Equation 1 reads 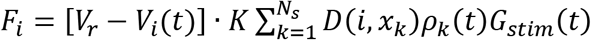, where *V_r_* = 20 mV is the excitatory reverse potential, and *K* the stimulation intensity.

### Macroscopic measurements

We measure the strength of the coupling within the neuronal population at time *t* by calculating their total synaptic weight (averaged over the neuron population) 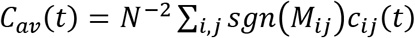, where *M_ij_* is defined in Equation 2, *sgn* is the sign-function, while *C_av_* is calculated by averaging over the last 100. *T_s_*. The extent of in-phase synchronization within the network is assessed by the order parameter (Haken, 1983;Kuramoto, 2012) *R*(*t*) = |*N*^−1^ Σ_*j*_*e*^*iφj*(*t*)^|, where *φ_j_*(*t*) = 2*π*(*t* – *t_j,m_*)/(*t*_*j,m*+1_ – *t_j,m_*) for *t_j,m_* ≤ *t* ≤ *t*_*j,m*+1_ is a linear approximation of the phase of neuron *j* between its *m^th^* and (*m* + 1)^*th*^ spikes at spiking times and *t*_*j,m*+1_. *R*(*t*) = 1 for complete in-phase synchronization, and *R*(*t*) = 0 in the absence of in-phase synchronization. Because of strong fluctuations of the order parameter, we calculate the moving average < *R* > over a time window of 400 · *T_s_*, to investigate the time evolution of the order parameter. Moreover, we use the quantity *R_av_*, which is the order parameter *R*(*t*) averaged over the last 100 · *T_s_* of a pause following a CR shot or of the end of the post-stim epoch. For the statistical description and analysis of the non-Gaussian distributed *R_av_* data (*n* = 11 samples), we use boxplots (Tukey, 1977). Their Inter-Quartile Range measures the statistical dispersion around the median, which is defined as width of the middle 50% of the distribution and is represented by a box. It is also used to determine outliers in the data: an outlier falls more than 1.5 times IQR below the 25% quartile or more than 1.5 times IQR above the 75% quartile.

### Dependence of CR stimulation outcome on CR stimulation frequency and intensity

This section provides a short overview of the results of a study where CR stimulation frequency and intensity were varied in detail (Manos et al., 2017). That study revealed the dependence of the outcome of RVS and SVS CR on the CR stimulation frequency and intensity and, in particular, possible limitations thereof, especially for SVS CR stimulation. Based on these limitations, the present study presents an approach that enables to overcome these issues.

In the present study, for each initial network condition and its corresponding parameters (simply denoted as *network*), we apply RVS and SVS CR stimulation with different realizations of the CR sequence orders per network. We start the simulations with an equilibration phase without STDP, which lasts for 2 s. From this point on, the network evolves in the presence of STDP, starting with a 60 s integration with STDP only (i.e. without stimulation), where a rewiring of the connections takes place, resulting in a strongly synchronized state with intrinsic firing rate f_int_ ≈ 71.4 Hz (corresponding to a period of *T*_int_ = 14 ms). We then run four different CR stimulation protocols, resetting the starting time to *t* = 0 s. We use 3:2 ON-OFF CR stimulation, where three stimulation ON-cycles (with stimulation on) alternated with two OFF-cycles (without stimulation), with ON-/OFF-cycle duration of *T_s_*. 3:2 ON-OFF CR stimulation was used in a number of computational, pre-clinical and clinical studies, for details and motivation see (Manos et al., 2017).

To study dosage regimens that potentially improve reliability and stimulation outcome of RVS and/or SVS stimulation, in the present study we focus on parameter ranges where RVS and/or SVS CR stimulation did not reliably elicit long-lasting desynchronization according to (Manos et al., 2017). In (Manos et al., 2017), we delivered CR single shots of 128 s duration followed by a 128 s CR-off period and varied the CR stimulation frequency and intensity over a wider range. In this way, we showed that RVS CR stimulation turned out to be more robust against variations of the stimulation frequency, while SVS CR stimulation can obtain stronger anti-kindling effects. This dependence on the CR stimulation intensity and frequency is summarized in **Figure 1. Figures 1A,1E** show the boxplots for the time-averaged mean synaptic weights *C_av_* (at the end of the 128 s CR-off period) with values belonging to the same intensity value *K* for RVS and SVS CR stimulation, respectively, but CR stimulation frequency varying in the interval [25%f_0_, …, 175%f_0_], with f_0_ as defined below. The motivation for restricting the CR stimulation intensity to the interval *K* ∈ [0.20,…, 0.50] is discussed in (Manos et al., 2017). In a similar manner, **Figures 1B,1F** show boxplots for the time-averaged order parameter *R_av_* (again at the end of the CR-off period) with values belonging to the same intensity value *K* for RVS and SVS CR stimulation respectively. **Figures 1C,1G** depict the boxplots for the time-averaged mean synaptic weights *C_av_* (at the end of the CR-off period) but now with values belonging to the same frequency ratio (f_stim_/f_0_) · 100 for RVS and SVS CR stimulation, respectively, but intensity value *K* ∈ [0.20,…, 0.50]. The CR stimulation frequency f_stim_ takes values in the interval [25%f_0_, …, 175%f_0_], where f_0_ = 1/*T*_0_ denotes the initial stimulation frequency. The choice of the frequency f_0_ (or period *T*_0_) of the CR stimulation is made with respect of the intrinsic network’s firing rate frequency (or period) f_int_(or *T*_int_) and is meant to have a value close to that (in this case *T*_0_ = 16 ms with f_0_ = 62.5 Hz). More details can be found in (Manos et al., 2017). **Figures 1D,1H** show the boxplots for the time-averaged order parameter *R_av_* (at the end of the CR-off period) with values belonging to the same frequency ratio (f_stim_/f_0_) · 100 for RVS and SVS CR stimulation, respectively.

**Figure 1.**
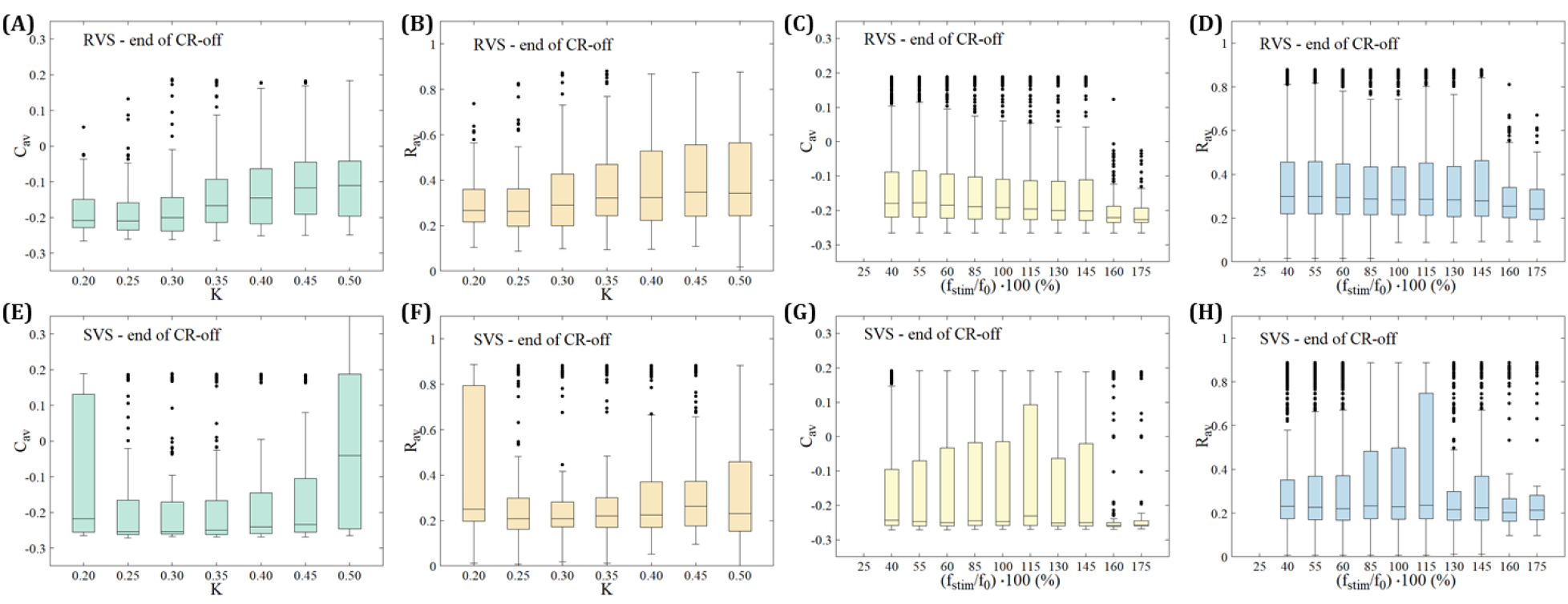
Dependence of stimulation outcome on CR stimulation intensity and frequency. **(A,E)** Boxplots for the time-averaged mean synaptic weights *C_av_* (at the end of the CR-off period) with values belonging to the same intensity value *K* for RVS (top row) and SVS CR (bottom row) stimulation, respectively. **(B,F)** Boxplots for the time-averaged order parameter *R_av_* (at the end of the CR-off period) with values belonging to the same intensity value *K* for RVS (top row) and SVS CR (bottom row) CR stimulation respectively. **(C,G)** Boxplots for the time-averaged mean synaptic weights *C_av_* (at the end of the CR-off period) with values belonging to the same frequency ratio (f_stim_/f_0_) · 100 for RVS (top row) and SVS CR (bottom row) CR stimulation respectively. **(D,H)** Boxplots for the time-averaged order parameter *R_av_* (at the end of the CR-off period) with values belonging to the same frequency ratio (f_stim_/f_0_) · 100 for RVS (top row) and SVS CR (bottom row) CR stimulation respectively.

## Results

### Simulation Description

We investigate two singleshot and two multishot, spaced CR stimulation protocols (**Figure 2**). The multishot *Protocols A* and *C* consist of five single CR shots of 128 s duration, each followed by a pause of 128 s, respectively (**Figures 2A,2C**). The CR singleshot *Protocol B* consists of a long singleshot of 5 × 128 s followed by a pause of 5 × 128 s (**Figures 2B**). The CR singleshot *Protocol D* consists of a long singleshot consisting of five single shots of 128 s duration, strung together without pauses in between, followed by a pause of 5 × 128 s (**Figures 2D**). The integral stimulation duration is identical for *Protocols A* and *B*. In *Protocols A* and *B* all stimulation parameters are kept constant. In contrast, in *Protocol C* at the end of each pause the amount of synchrony is evaluated in a time window of 100 stimulation periods length (**Figure 2**) and a three-stage control scheme is put in place: (i) If the amount of synchrony does not fall below a pre-defined threshold, the CR stimulation frequency is mildly varied. (ii) If the desynchronization effect is moderate, the CR stimulation frequency remains unchanged. (iii) If desynchronization is achieved, the stimulation intensity is set to zero for the subsequent shot. Analogously, in *Protocol D* at the end of each single shot the amount of synchrony is evaluated in a time window of 100 stimulation periods length (**Figure 2**) and the three-stage control scheme is executed. The difference between *Protocol C* and *D* is that the evaluation for the control intervention is performed in a pause subsequent to a single shot (*Protocol C*) as opposed to during a single shot (*Protocol D*).

**Figure 2.**
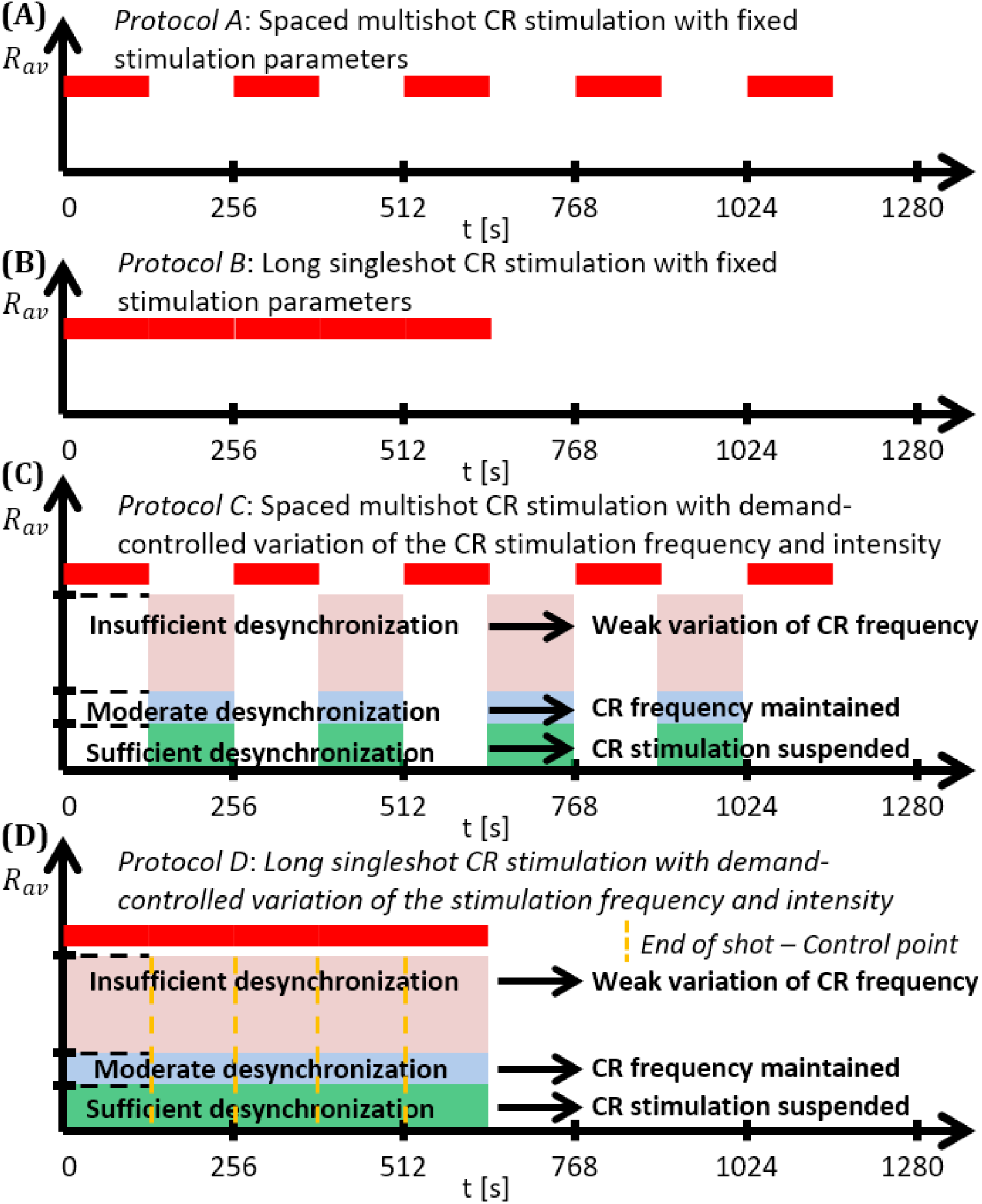
Schematic summary of the CR stimulation protocols. **(A)** *Protocol A*: Spaced multishot CR stimulation with fixed stimulation parameter. **(B)** *Protocol B*: Long singleshot CR stimulation with fixed stimulation parameters. **(C)** *Protocol C*: Spaced multishot CR stimulation with demand-controlled variation of the CR stimulation frequency and intensity. **(D)** *Protocol D*: Long singleshot CR stimulation with demand-controlled variation of the stimulation frequency and intensity (see text).

For the stage (i) control, the variation of the CR stimulation frequency is not adapted to frequency characteristics of the neuronal network. Rather a minor variation of the CR stimulation frequency is performed to make a fresh start with the subsequent single CR shot. These minor changes of the CR stimulation frequency do not lead to changes of the neurons’ intrinsic firing rates of more than ±3%.

Due to the stage (iii) control, the demand-controlled shutdown of CR stimulation, the maximum integral stimulation duration of *Protocol C* and *D* can reach the level of *Protocols A* and *B*, but may well fall below. We use the order parameter to assess the amount of synchronization (see *Materials and Methods*).

#### Protocol A: Spaced multishot CR stimulation with fixed stimulation parameters

For this stimulation protocol all stimulation parameters are kept constant (**Figure 2**). Accordingly, the CR stimulation period *T_s_* remains constant, too. We study the stimulation outcome of only five symmetrically spaced consecutive single CR shots. To this end, for both RVS CR and SVS CR stimulation we consider two unfavorable parameter pairs of fixed CR stimulation period and intensity, respectively. One example refers to cases where CR stimulation induces acute effects, but no long-lasting desynchronizing effects (*Cases I and IV*). The other example concerns the case where CR stimulation causes neither acute nor long-lasting desynchronizing effects in a reliable manner (*Cases II and III*).

##### RVS CR stimulation: Case I

(*K,T_s_*) = (0.30,11). At a stimulation duration of 128 s these parameters caused only an acute, but no long-lasting desynchronization in the majority of networks studied [**Figure 4B** of (Manos et al., 2017), where *T_s_* = 11 ms corresponds to ~127% of the intrinsic firing rate (or ~91 Hz)]. *Case II*: (*K,T_s_*) = (0.20,28). In the majority of networks tested, these parameters did neither lead to acute nor long-lasting desynchronization after administration of a single CR shot [**Figure 5B** of (Manos et al., 2017) where *T_s_* = 28 ms corresponds to ~50% of the intrinsic firing rate (or ~36 Hz)]. For both cases, we investigate the order parameter < *R* > averaged over a sliding window for 11 different networks (marked with different color/line types) (**Figures 2A,2C**). Boxplots of the order parameter *R_av_* averaged over a window of length 100 · *T_s_* at the end of each pause demonstrate the overall stimulation outcome for all tested 11 networks (**Figures 2B,2D**).

##### Case I

RVS CR stimulation induces a desynchronization during the CR shots (**Figure 3A**), but no reliable, long-lasting desynchronization in the subsequent pauses (**Figure 3B**). The spacing protocol with five identical RVS CR shots does not significantly improve the desynchronizing outcome of a single RVS CR shot. In fact, in the boxplots the large dispersion around the median value remains almost unchanged in the course of this protocol (**Figure 3B**). *Case II*: Neither during the RVS CR shots nor during the subsequent pauses a sufficient desynchronization is observed (**Figures 2C,2D**). The spacing protocol does not cause an improvement of the stimulation outcome in this case, too (**Figure 3D**).

**Figure 3.**
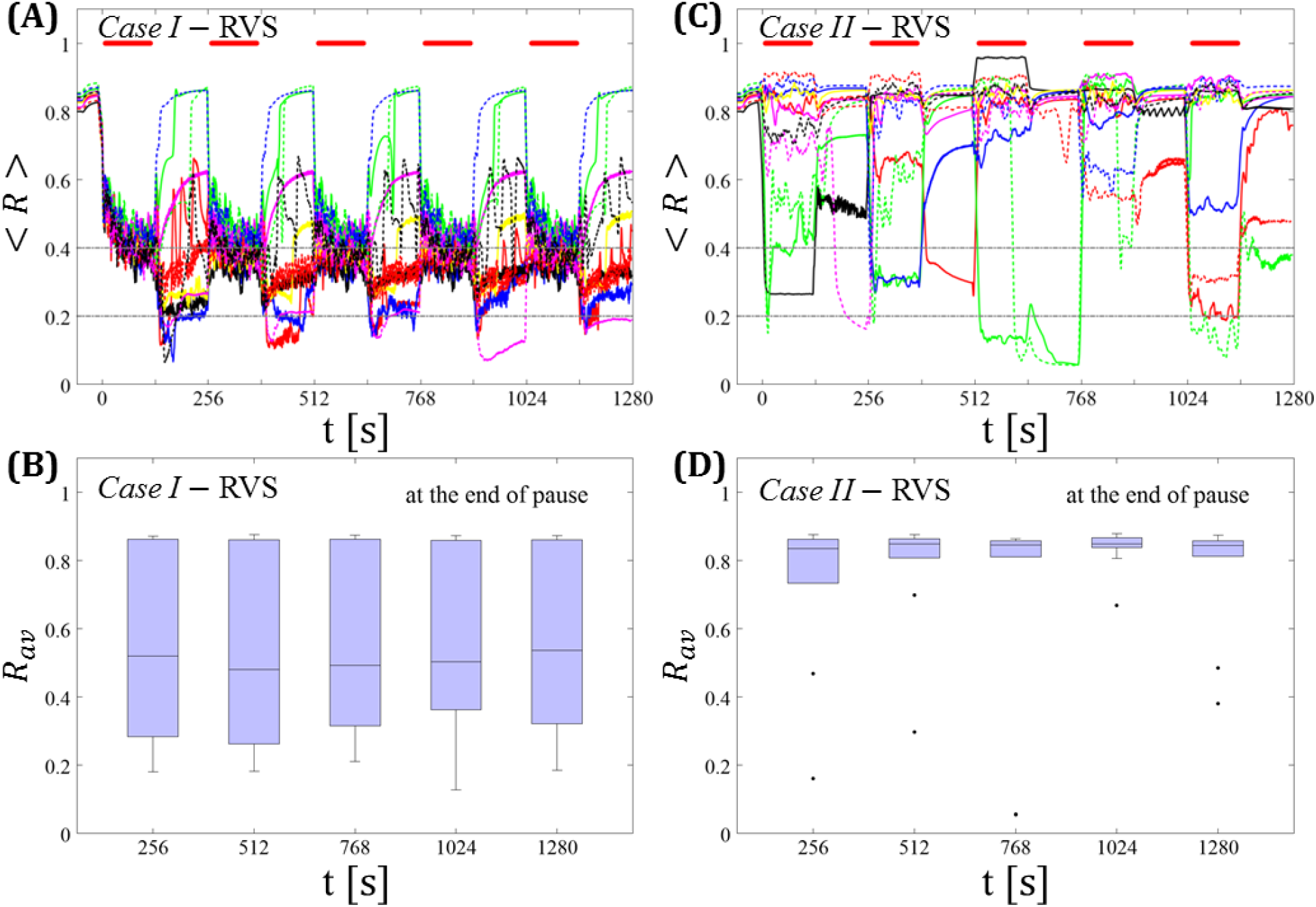
Protocol A: Spaced multishot RVS CR stimulation with fixed stimulation period *T_s_*. **(A,C)** Time evolution of the order parameter < *R* > averaged over a sliding window during 5 consecutive RVS CR shots with fixed CR stimulation period. Different colors correspond to different networks. Stimulation parameters are unfavorable for anti-kindling in *Case I* **(A,B)** and *Case II* **(C,D)** (see text). **(A,C)** The horizontal solid red lines indicate the CR shots, while the horizontal dashed grey lines serve as visual cues. Spacing is symmetrical, i.e. CR shots and consecutive pauses are of the same duration. **(B,D)** Boxplots for *R_av_*, averaged over a window of length 100 · *T_s_* at the end of each pause, illustrate the overall outcome for all tested 11 networks. *Case I*: (*K, T_s_*) = (0.30,11). *Case II*: (*K,T_s_*) = (0.20,28).

##### SVS CR stimulation: Case III

(*K, T_s_*) = (0.20,9). Single shot SVS CR stimulation with these parameters caused neither pronounced acute nor long-lasting desynchronization [**Figure 6D** of (Manos et al., 2017), where *T_s_* = 9 ms corresponds to ~156% of the intrinsic firing rate (or ~111 Hz)]. *Case IV*: (*K,T_s_*) = (0.20,14). Single shot SVS CR stimulation with these parameters led to acute, but no long-lasting desynchronization in the majority of networks tested [**Figure 7B** of (Manos et al., 2017) where *T_s_* = 28 ms corresponds to ~50% of the intrinsic firing rate (or ~36 Hz)]. For both cases we performed the same analysis as shown in **Figure 3**.

##### Case III

SVS CR stimulation neither induces a pronounced and reliable desynchronization during the CR shots (**Figure 4A**) nor during the pauses (**Figure 4B**). In fact, the dispersion around the median value is increased during the fourth and fifth pause (**Figure 4B**).

**Figure 4.**
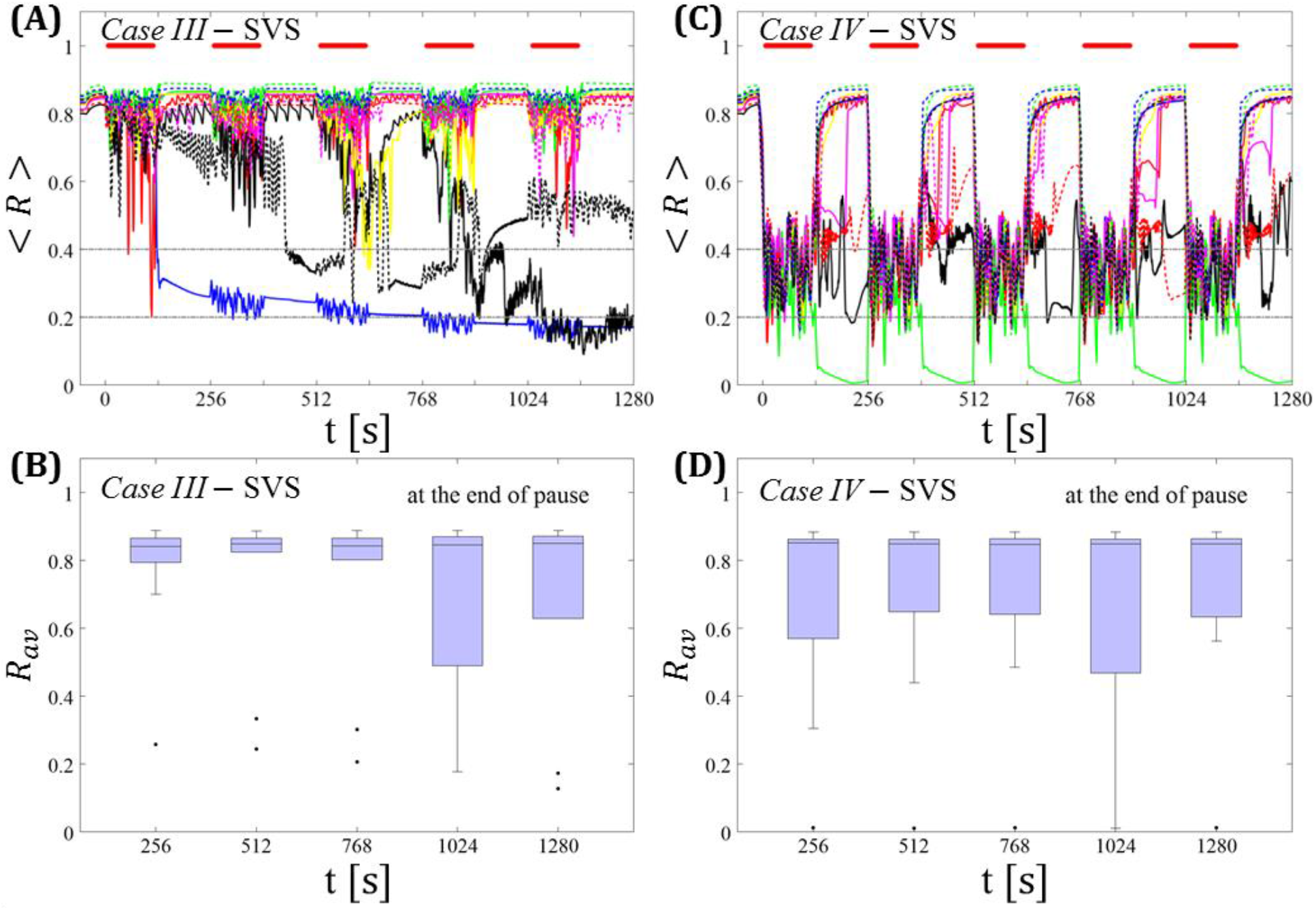
Protocol A: Spaced multishot SVS CR stimulation with fixed stimulation period *T_s_*. **(A,C)** Time evolution of the time-averaged order parameter < *R* > during 5 consecutive SVS CR shots with fixed CR stimulation period. Stimulation parameters are unfavorable for anti-kindling in *Case III* **(A,B)** and *Case IV* **(C,D)** (see text). **(B,D)** Boxplots for *R_av_*, averaged over a window of length 100 · *T_s_* at the end of each pause, illustrate the overall outcome for all tested 11 networks. *Case III*: (*K, T_s_*) = (0.20,9). *Case IV*: (*K, T_s_*) = (0.20,14). Same format as in **Figure 3**.

##### Case IV

During the SVS CR shots a desynchronization occurs (**Figure 4C**). However, no reliable and pronounced desynchronization is observed during the pauses (**Figure 4D**).

In summary, for both RVS CR and SVS CR stimulation the spacing protocol with five consecutive CR shots does not cause an improvement of the long-lasting desynchronization (assessed after cessation of stimulation). We performed the same analysis for a larger set of (*K, T_s_*) pairs in the parameter plane analyzed in (Manos et al., 2017), with *K* ranging from 0.2 to 0.3 (weak intensities) and *T_s_* ranging from 9 ms to 28 ms (around the intrinsic period). For all parameter pairs tested, the spacing *Protocol A* did not improve the long-term desynchronization effect.

#### Protocol B: Long singleshot CR stimulation with fixed stimulation parameters

For this stimulation protocol all stimulation parameters are kept constant, too (**Figure 2**). Instead of five single CR shots of 128 s duration each (**Figures 1–3**), we deliver one fivefold longer singleshot of 5 × 128 s duration (**Figure 2**). For both RVS CR and SVS CR stimulation we consider the corresponding two unfavorable parameter pairs of fixed CR stimulation period and intensity already studied above (*Cases I-IV*). This is to study whether a fivefold prolongation of the stimulation duration leads to an improvement of the stimulation outcome.

##### RVS CR stimulation

For comparison, we consider the cases studied above. *Case I*: (*K, T_s_*) = (0.30,11). *Case II*: (*K,T_s_*) = (0.20,28). We study the order parameter < *R* > averaged over a sliding window for 11 different networks (marked with different color/line types in **Figure 5A**). The overall stimulation outcome for all tested 11 networks is illustrated with boxplots of the order parameter *R_av_* averaged over a window of length 100 · *T_s_* at the end of the post-stimulus epoch (**Figure 5B**).

**Figure 5.**
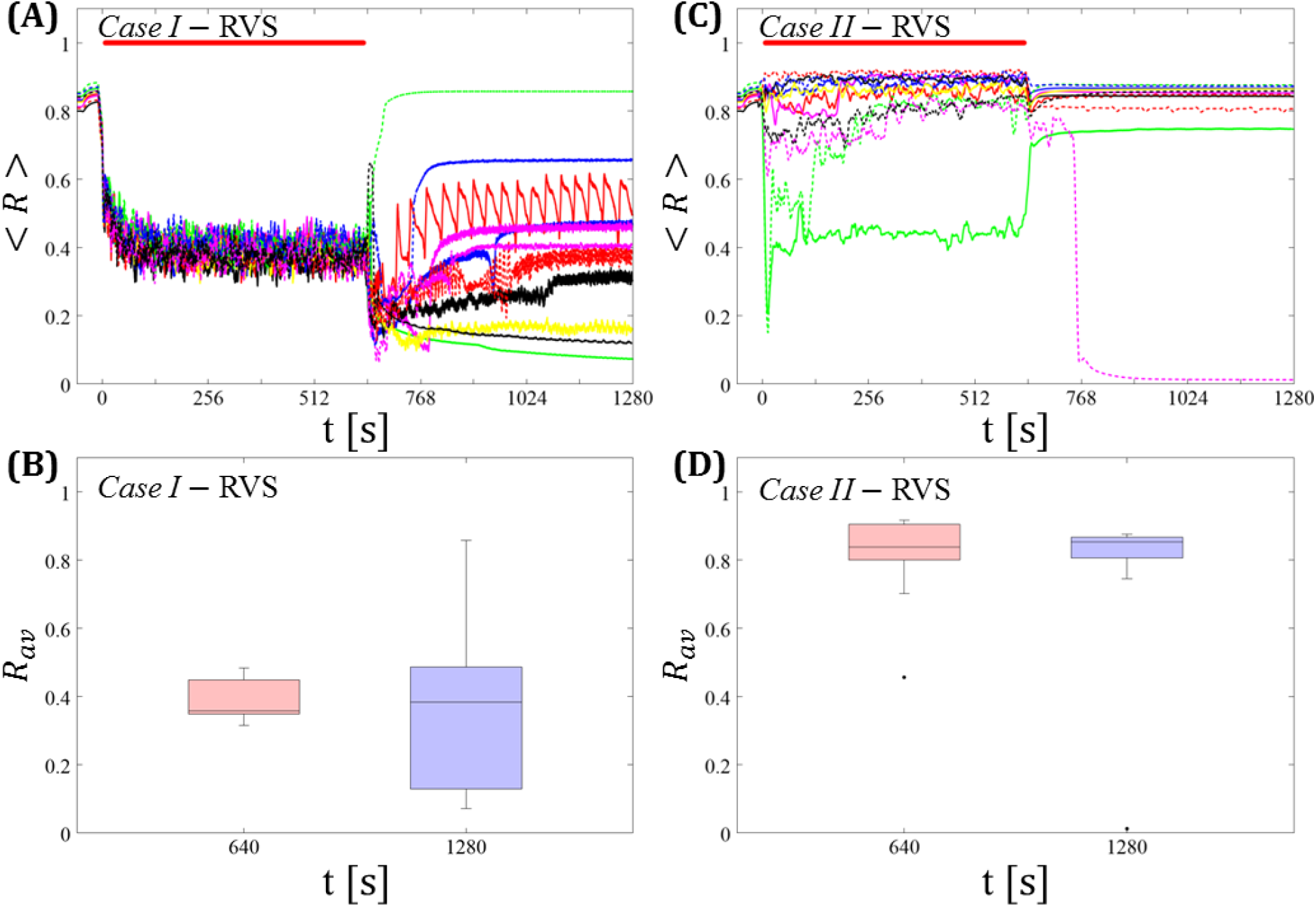
Protocol B: Long singleshot RVS CR stimulation with fixed *T_s_*. **(A,C)** Time evolution of the time-averaged order parameter < *R* > during and after one long RVS CR singleshot with fixed CR stimulation period. Stimulation parameters are unfavorable for anti-kindling for short singlehots of 128 s duration in *Case I* **(A,B)** and *Case II* **(C,D)** (see text). **(B,D)** Boxplots for *R_av_*, averaged over a window of length 100 · *T_s_* at the end of the singleshot and at the end of the post-stim epoch illustrate the overall outcome for all tested 11 networks. *Case I*: (*K,T_s_*) = (0.30,11). *Case II*: (*K, T_s_*) = (0.20,28). Same format as in **Figure 3**.

##### Case I

RVS CR stimulation induces a desynchronization during the long RVS CR singleshot (**Figure 5A**), but no reliable, long-lasting desynchronization in the subsequent pauses (**Figure 5B**). The median of the order parameter of the long-term outcome hardly changes, but the dispersion is significantly greater for the post-stim order parameter (**Figure 5B**). Note, the overall long-term desynchronization for the long singleshot (**Figure 5B**) is more pronounced compared to the spaced RVS CR stimulation *Protocol A* (**Figure 3B**).

##### Case II

Neither during the long RVS CR singleshot nor during the subsequent stimulation-free epoch a reliable and pronounced desynchronization is observed (**Figures 4C,4D**). Interestingly, the network that undergoes an acute desynchronization during the singleshot relaxes back to a synchronized state (**Figure 5B**, green curve). Conversely, the only network that displays a long-term desynchronization does not undergo a pronounced desynchronization during the singleshot (**Figure 5B**, magenta curve).

##### SVS CR stimulation

For comparison, we consider the time course of the time-averaged order parameter < *R* > (**Figures 5A,5C**) and the corresponding boxplots of the order parameter *R_av_* averaged over a window of length 100 · *T_s_* at the end of the post-stimulus epoch (**Figures 5B,5D**) for 11 different networks (marked with different color/line types) for the cases studied above. *Case III*: (*K,T_s_*) = (0.20,9). *Case IV*: (*K,T_s_*) = (0.20,14).

##### Case III

In the majority of networks SVS CR stimulation does not induce a pronounced and reliable desynchronization during the fivefold longer SVS CR singleshot as well as in the post-stim epoch (**Figure 6A**). Three out of 11 networks display a pronounced acute and long-lasting desynchronization (**Figure 6A**). Accordingly, the dispersion around the median is large during and after the singleshot (**Figure 6B**).

**Figure 6.**
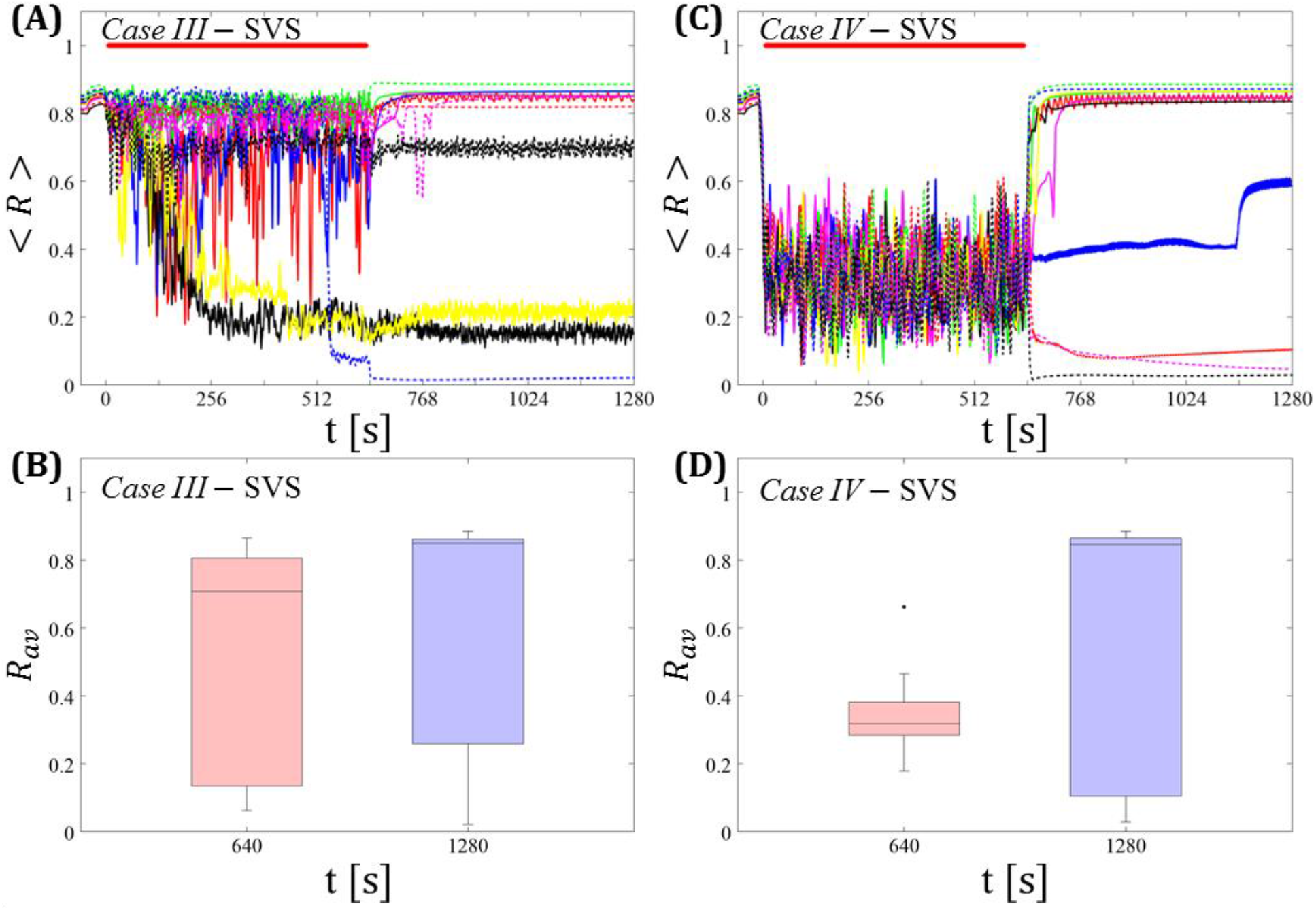
Protocol B: Long singleshot SVS CR stimulation with fixed *T_s_*. **(A,C)** Time evolution of the time-averaged order parameter < *R* > during and after one long SVS CR singleshot with fixed CR stimulation period. Stimulation parameters are unfavorable for anti-kindling for short singlehots of 128 s duration in *Case III* **(A,B)** and *Case IV* **(C,D)** (see text). **(B,D)** Boxplots for *R_av_*, averaged over a window of length 100 · *T_s_* at the end of the singleshot and at the end of the post-stim epoch illustrate the overall outcome for all tested 11 networks. *Case III*: (*K, T_s_*) = (0.20,9). *Case IV*: (*K, T_s_*) = (0.20,14). Same format as in **Figure 3**.

##### Case IV

During the long SVS CR singleshot a pronounced desynchronization occurs (**Figure 6C**), as reflected by the small dispersion around the small median in the corresponding boxplot (**Figure 6D**). However, in the post-stimulation epoch most of the networks relax to a synchronized state, with only a few networks remaining in a long-term desynchronized state (**Figure 6D**). Accordingly, there is a large dispersion around a large median in the boxplot (**Figure 6D**).

In summary, for both RVS CR and SVS CR stimulation the fivefold increase of the stimulation duration does not lead to a reliable and pronounced long-lasting desynchronization. Again, we performed the same analysis for a larger set of (*K, T_s_*) pairs in the parameter plane analyzed in (Manos et al., 2017), with *K* ranging from 0.2 to 0.3 (weak intensities) and *T_s_* ranging from 9 ms to 28 ms (around the intrinsic period value). For all parameter pairs tested, the spacing *Protocol B* did not lead to a reliable and pronounced long-term desynchronization.

#### Protocol C: Spaced multishot CR stimulation with demand-controlled variation of stimulation period *T_s_* and intensity

We study the stimulation outcome of only five symmetrically spaced consecutive single CR shots with stimulation period *T_s_* and intensity varied according to a three-stage control scheme. To this end, for both RVS CR and SVS CR stimulation we consider two unfavorable parameter pairs of fixed CR stimulation period and intensity, respectively. One example refers to cases where CR stimulation induces acute effects, but no long-lasting desynchronizing effects (*Cases I* and *IV*). The other example concerns the case where CR stimulation causes neither acute nor long-lasting desynchronizing effects in a reliable manner (*Cases II* and *III*). We consider a regular and a random type of demand-controlled variation of the CR stimulation period *T_s_*. Note, in both cases the CR stimulation period is not adapted to frequency characteristics of the network. We consider the time courses of the time-averaged order parameter < *R* > and *R_av_*, the order parameter averaged over a window of length 100 · *T_s_* at the end of pause.

##### Demand-controlled regular variation of the CR stimulation period and demand-controlled variation of the intensity

At the end of each pause we calculate the order parameter *R_av_* averaged over a window of length 100 · *T_s_*. We vary the CR stimulation period and intensity according to the amount of synchrony, based on a three-stage control scheme:

i. *Insufficient desynchronization*: If *R_av_* > 0.4, we decrease the CR stimulation period of the subsequent RVS shot by *T_s_*(*j* + 1) = *T_s_*(*j*) −1 ms, where the index *j* stands for the *j*-th CR shot. As lower bound we set *T_s_* = 9 ms (corresponding to ~156% of the intrinsic firing rate), in order to avoid undesirably high CR stimulation frequencies. In a previous computational study the latter turned out to be unfavorable for desynchronization [see (Manos et al., 2017)]. As soon as *T_s_* reaches its lower bound of 9 ms, it is reset to *T_s_*(1) + 1 ms.
ii. *Moderate desynchronization*: If 0.2 ≤ *R_av_* ≤ 0.4, we preserve the CR stimulation period for the subsequent CR shot: *T_s_*(*j* + 1) = *T_s_*(*j*), where the index *j* denotes the *j*-th CR shot. 0.2 ≤ *R_av_* ≤ 0.4 is considered to be indicative of a desynchronization effect.
iii. *Sufficient desynchronization*: If *R_av_* < 0.2, the CR stimulation is suspended for the subsequent shot by setting *K* = 0 for the next shot and until 0.2 ≤ *R_av_*. *R_av_* ≤ 0.2 is considered a sufficient desynchronization.

##### Spaced multishot RVS CR stimulation with demand-controlled regular variation of the stimulation period T_s_ and demand-controlled variation of the intensity

In both *Cases* (*I* and *II*) this protocol reliably induces a desynchronization for all networks tested (**Figures 6A,6C**). After the second RVS CR shot the median of the time-averaged order parameter *R_av_* at the end of the corresponding pauses falls below 0.4, with moderate dispersion (**Figures 6B,6D**). Note, already after the first mild variation of the CR stimulation period *T_s_* the amount of synchrony is significantly reduced. In several networks and pauses, the desynchronization criterion, *R_av_* < 0.2, is fulfilled, so that during the subsequent CR shots no stimulation is delivered (**Figures 6A,6C**). Accordingly, *Protocol C* enables to reduce the integral amount of stimulation.

**Figure 7.**
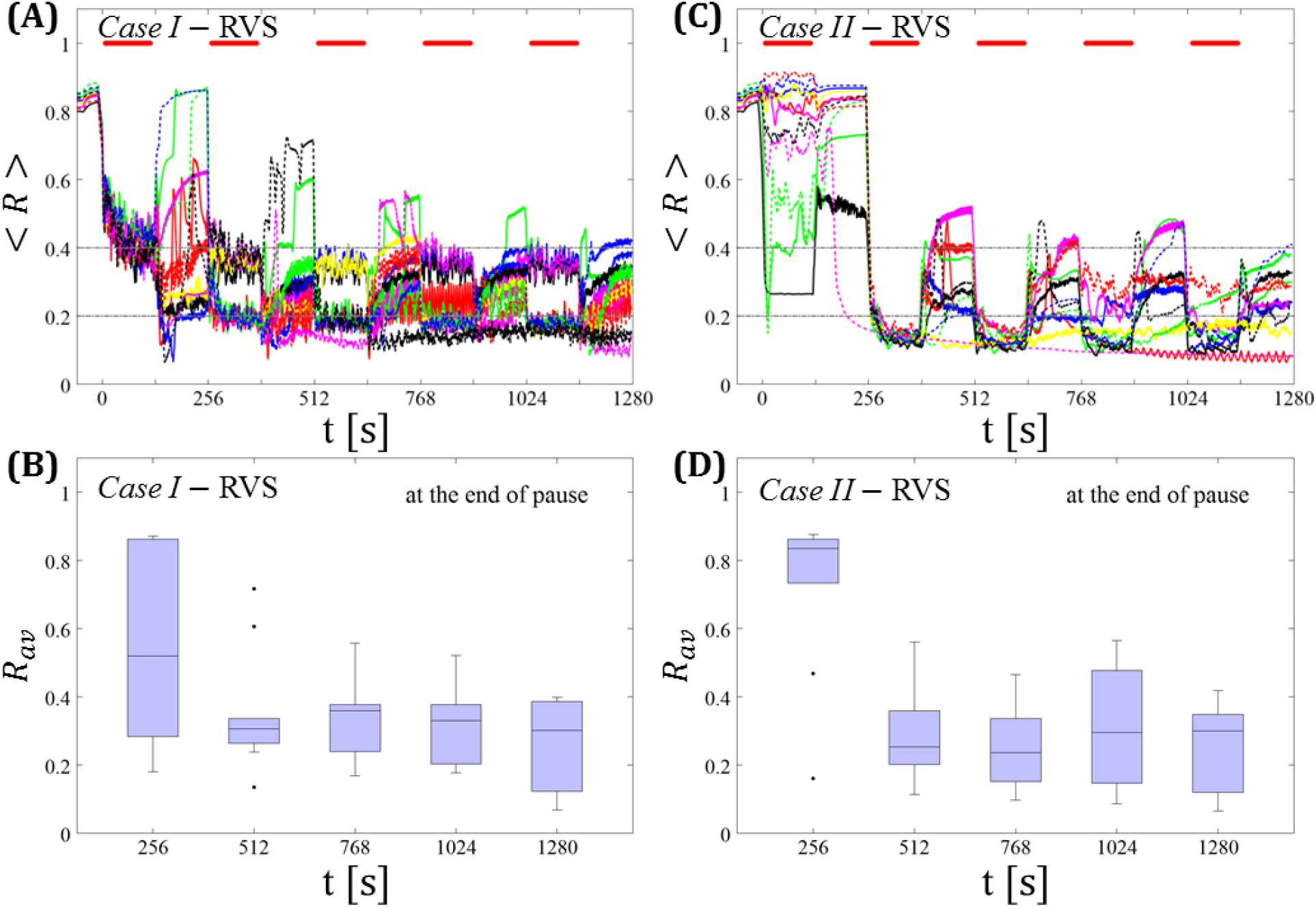
Protocol C: Spaced multishot RVS CR stimulation with demand-controlled regular variation of the stimulation period *T_s_* and with demand-controlled variation of the intensity. **(A,C)** Time evolution of the order parameter < *R* > averaged over a sliding window during 5 consecutive RVS CR shots. If *R_av_*, the order parameter averaged over a window of length 100 · *T_s_* at the end of a pause, exceeds 0.4, the CR stimulation period of the subsequent RVS shot is decreased by *T_s_* → *T_s_* — 1 ms (see text). Stimulation parameters are unfavorable for anti-kindling in *Case I* **(A,B)** and *Case II* **(C,D)** (see text). **(A,C)** The horizontal solid red lines indicate the CR shots, while the horizontal dashed grey lines serve as visual cues. Spacing is symmetrical, i.e. CR shots and consecutive pauses are of the same duration. **(B,D)** Boxplots for the time-averaged order parameter *R_av_* at the end of each pause, illustrate the overall outcome for all tested 11 networks. *Case* I: (*K,T_s_*) = (0.30,11). *Case II*: (*K,T_s_*) = (0.20,28).

##### Spaced multishot SVS CR stimulation with demand-controlled regular variation of the stimulation period T_s_ and demand-controlled variation of the intensity

This protocol causes a desynchronization for all networks tested in both *Cases* (*III* and *IV*) (**Figures 7A,7C**). After the third (*Case III*, **Figure 7B**) or the second SVS CR shot (*Case IV*, **Figure 7D**) a pronounced desynchronization is achieved, as reflected by a median of *R_av_* close to 0.2 (**Figures 7B,7D**). Accordingly, about half of the networks fulfilled the desynchronization criterion *R_av_* < 0.2 after the third SVS CR shot and, hence, did not require further CR stimulation. The mean firing rate, measured at the end of each pause did not deviate from the baseline firing rates by more than ±3%, irrespective of the extent of protocol-induced variation of the stimulation period *T_s_* (data not shown).

**Figure 8.**
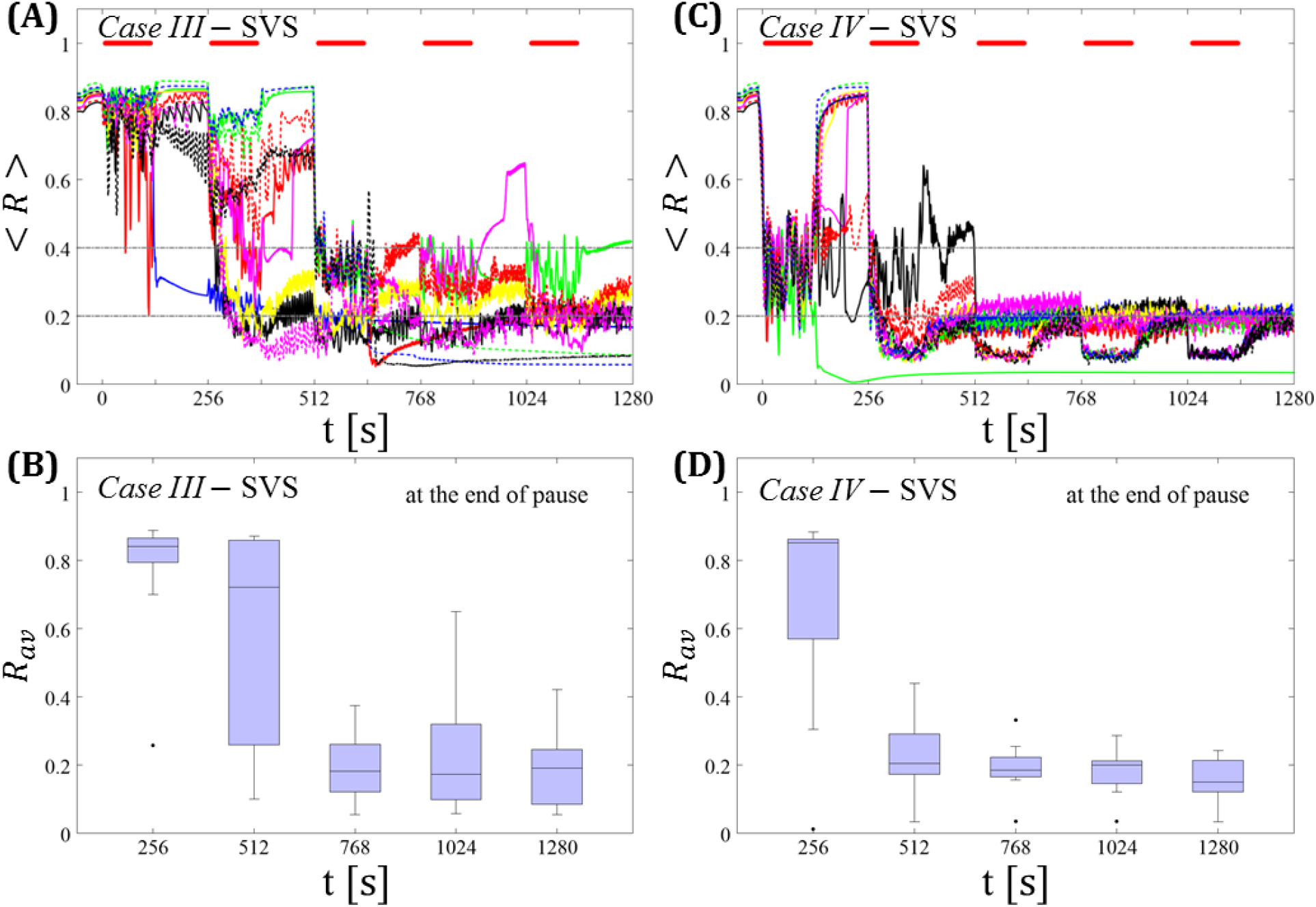
Protocol C: Spaced multishot SVS CR stimulation with demand-controlled regular variation of the stimulation period *T_s_* and with demand-controlled variation of the intensity. **(A,C)** Time evolution of the order parameter < *R* > averaged over a sliding window during 5 consecutive SVS CR shots. If *R_av_*, the order parameter averaged over a window of length 100 · *T_s_* at the end of a pause, exceeds 0.4, the CR stimulation period of the subsequent SVS shot is decreased by *T_s_* → *T_s_* – 1 ms (see text). Stimulation parameters are unfavorable for anti-kindling in *Case III* **(A,B)** and *Case IV* **(C,D)** (see text). **(B,D)** Boxplots for the time-averaged order parameter *R_av_* at the end of each pause, illustrate the overall outcome for all tested 11 networks. *Case III*: (*K, T_s_*) = (0.20,9). *Case IV*: (*K, T_s_*) = (0.20,14). Same format as in **Figure 7**.

We do not adapt the CR stimulation period *T_s_* to frequency characteristics of the stimulated network. To further illustrate this aspect, we replace a regular, increasing or decreasing variation of the stimulation period by a random variation.

##### Demand-controlled random variation of the CR stimulation period and demand-controlled variation of the intensity

Again, at the end of each pause we calculate the order parameter *R_av_* averaged over a window of length 100 · *T_s_*. A random variation of the CR stimulation period is performed, depending on the amount of synchrony detected. To this end, we select the interval [*T_s_*(1) − 4 ms, *T_s_*(1) + 4 ms], where *T_s_*(1) denotes the CR stimulation period of the first shot. By design, this interval has a lower bound at 9 ms. The three-stage control scheme is governed by:

i. *Insufficient desynchronization*: If *R_av_* > 0.4 at the end of the pause of the *j*-th CR shot, we randomly pick *T_s_*(*j* + 1) and skip inefficient values used before.
ii. *Moderate desynchronization*: If 0.2 ≤ *R_av_* ≤ 0.4, we preserve the CR stimulation period for the subsequent CR shot: *T_s_*(*j* + 1) = *T_s_*(*j*), where the index *j* denotes the *j*-th CR shot.
iii. *Sufficient desynchronization*: If *R_av_* < 0.2, the CR stimulation is suspended for the subsequent shot by setting *K* = 0 for the next shot and until 0.2 ≤ *R_av_*.

The feasibility of this protocol is demonstrated by considering one example for RVS CR stimulation (*Case II*, **Figure 7**) and one for SVS CR stimulation (*Case IV*, **Figure 9**). For both cases we additionally provide the mean firing rate of the networks at the end of each shot and at the end of each subsequent pause to demonstrate that deviations do not exceed ±3% (**Figures 8C, 9C**). In the RVS case (**Figure 9**), the time course of the order parameter < *R* > (**Figure 9A**) and the corresponding boxplots of *R_av_* (**Figure 9B**) display a similar pattern of reliable desynchronization as obtained by *Protocol C* with regular variation of the CR stimulation duration *T_s_* (**Figure 7**). In principle, the SVS case (**Figure 10**) provides similar findings as with a regular variation of the CR stimulation duration *T_s_* (**Figure 7**). However, one network relaxes back to a strongly synchronized state (**Figure 10A**, dashed blue line). Delivering a sixth SVS CR shot with randomly varied *T_s_* to that network caused a desynchronization (data not shown). This example illustrates that a sequence of five SVS CR shots might not be sufficient to induce desynchronization in all possible networks.

**Figure 9.**
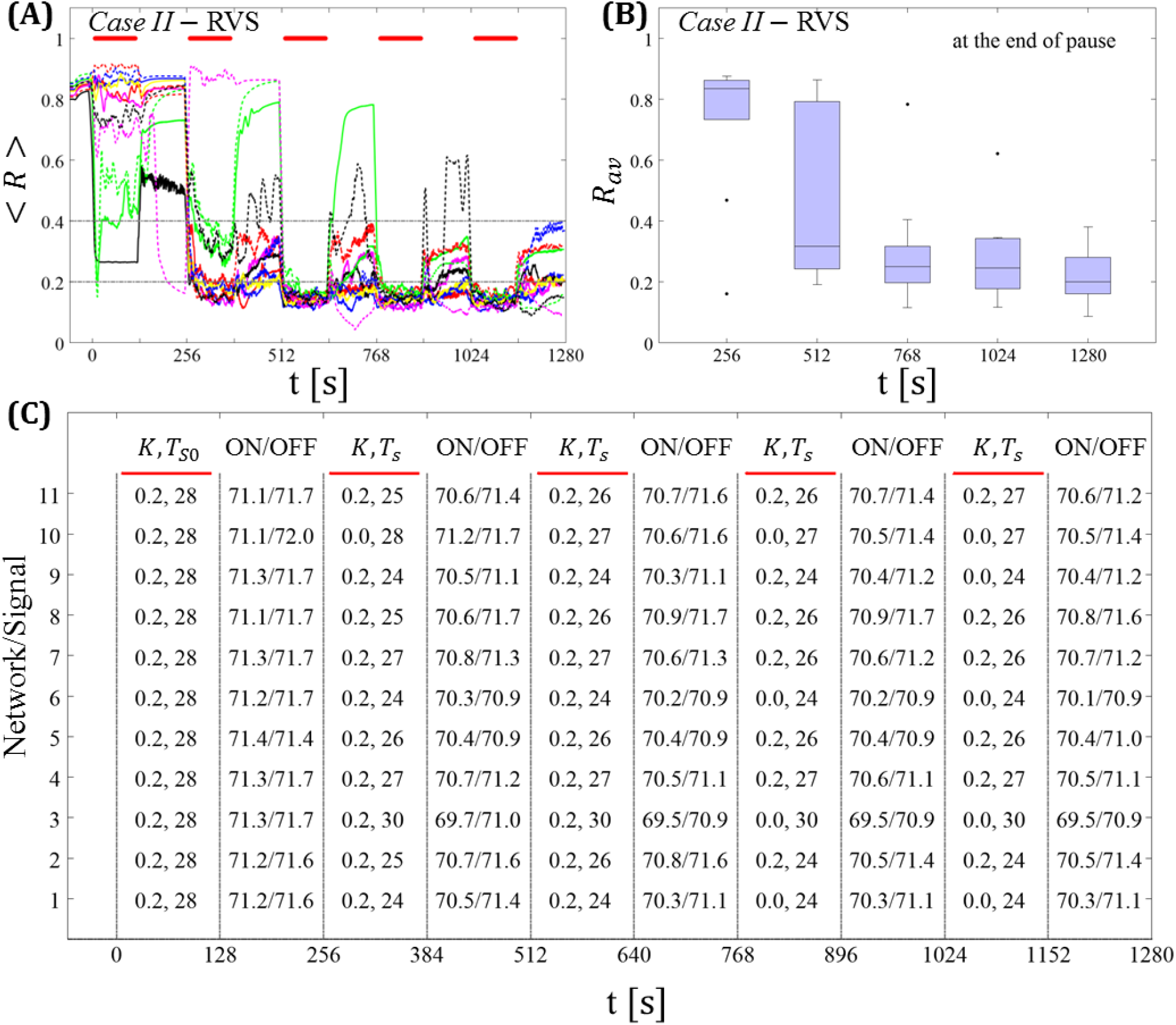
Protocol C: Spaced multishot RVS CR stimulation with demand-controlled random variation of the stimulation period *T_s_* and with demand-controlled variation of the intensity. **(A)** Time evolution of the order parameter < *R* > averaged over a sliding window during 5 consecutive RVS CR shots. The CR stimulation period is randomly varied depending on *R_av_*, by randomly picking a value from a narrow interval around the start period (see text). The horizontal solid red lines indicate the CR shots, while the horizontal dashed grey lines highlight the two control thresholds (see text). **(B)** Boxplots for the time-averaged order parameter *R_av_* at the end of each pause, illustrate the overall outcome for all tested 11 networks. **(C)** For each (vertically aligned) network the table presents CR intensity (*K* = 0.2 if CR is ON or *K* = 0 if CR is OFF during a CR shot) and stimulation period *T_s_* used for each CR shot (indicated by red bars) together with the mean firing rate of the network at the end of each CR shot (“ON”) and at the end of the subsequent pause (“OFF”). The mean firing rate was strongly fluctuating and, hence, calculated in a window of length 200 · *T_s_*. *Case II* stimulation parameters are unfavorable for anti-kindling: (*K, T_s_*) = (0.20,28) (see text).

**Figure 10.**
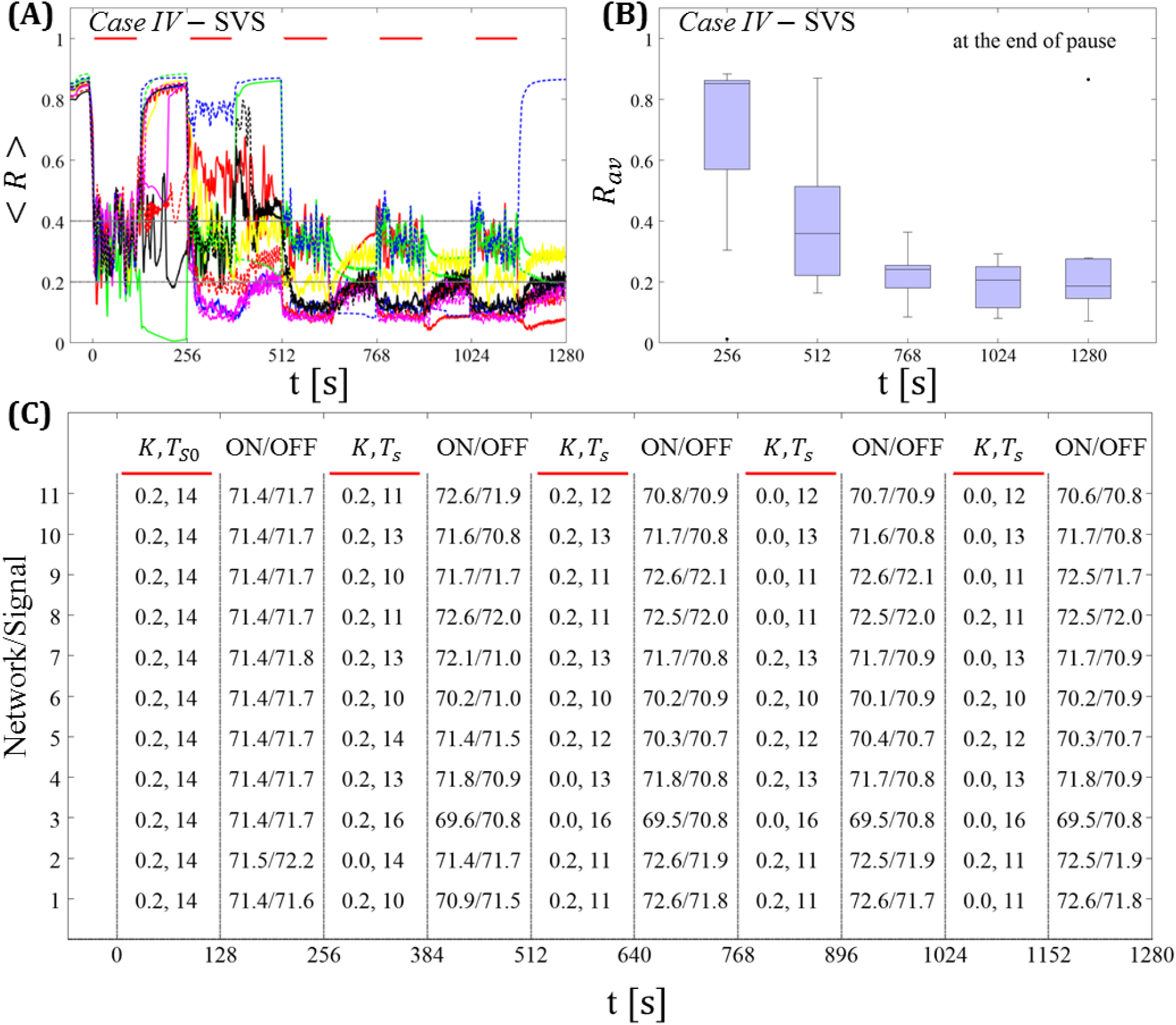
Protocol C: Spaced multishot SVS CR stimulation with demand-controlled random variation of the stimulation period *T_s_* and with demand-controlled variation of the intensity. **(A)** Time evolution of the order parameter < *R* > averaged over a sliding window during 5 consecutive RVS CR shots with demand-controlled random variation of *T_s_* (see text). **(B)** Boxplots for the time-averaged order parameter *R_av_* at the end of each pause, illustrate the overall outcome for all tested 11 networks. **(C)** For each (vertically aligned) network the table presents CR intensity (fr = 0.2 if CR is ON or *K* = 0 if CR is OFF during a CR shot) and stimulation period *T_s_* used for each CR shot (indicated by red bars) and the mean firing rate of the network at the end of each CR shot (“ON”) and at the end of the subsequent pause (“OFF”). *Case IV* stimulation parameters are unfavorable for anti-kindling: (*K, T_s_*) = (0.20,14) (see text). Same format as in **Figure 9**.

In summary, for the five-shot RVS CR as well as SVS CR stimulation *Protocol C* with regular as well as random variation of the CR stimulation duration *T_s_* we observed a pronounced desynchronization, with the exception of one network (Figure 10A, dashed blue line). Our analysis was performed for a larger set of (*K, T_s_*) pairs in the parameter plane analyzed in (Manos et al., 2017), with *K* ranging from 0.2 to 0.3 (weak intensities) and *T_s_* ranging from 9 ms to 28 ms (around the intrinsic period). For all parameter pairs tested, the spacing *Protocol C* with regular and random variation of *T_s_* led to a reliable and pronounced long-term desynchronization in the vast majority of networks tested.

#### Protocol D: Long singleshot CR stimulation with demand-controlled variation of the stimulation frequency

*Protocol D* consists of five consecutive shots. Unlike in *Protocol C*, there are no pauses between the five consecutive shots, so that they form one long singleshot.

##### Demand-controlled regular variation of the CR stimulation period and demand-controlled variation of the intensity

At the end of each shot we calculate the order parameter *R_av_* averaged over a window of length 100 · *T_s_*. We vary the CR stimulation period and intensity according to the amount of synchrony, based on the three-stage control scheme as used for *Protocol C* (see above).

##### Long singleshot RVS CR stimulation with demand-controlled variation of the stimulation frequency

In *Case I* this protocol seems to perform similarly well (**Figures 11A,11B**) as *Protocol C* (**Figures 7A,7B**) and *Protocol B* (**Figures 5A,5B**) which is also an alternative long singleshot but with fixed *T_s_*. After the second RVS CR shot almost all networks reach a *moderate* or *sufficient desynchronization* which is maintained fairly well after the RVS CR is ceased. Nonetheless, this particular protocol does not perform equally well for *Case II* (**Figures 11C,11D**). Even in the cases, where the variation of *T_s_* leads to some improvement, the overall long-lasting effect is worse than with *Protocol C* (**Figures 7C,7D**).

**Figure 11.**
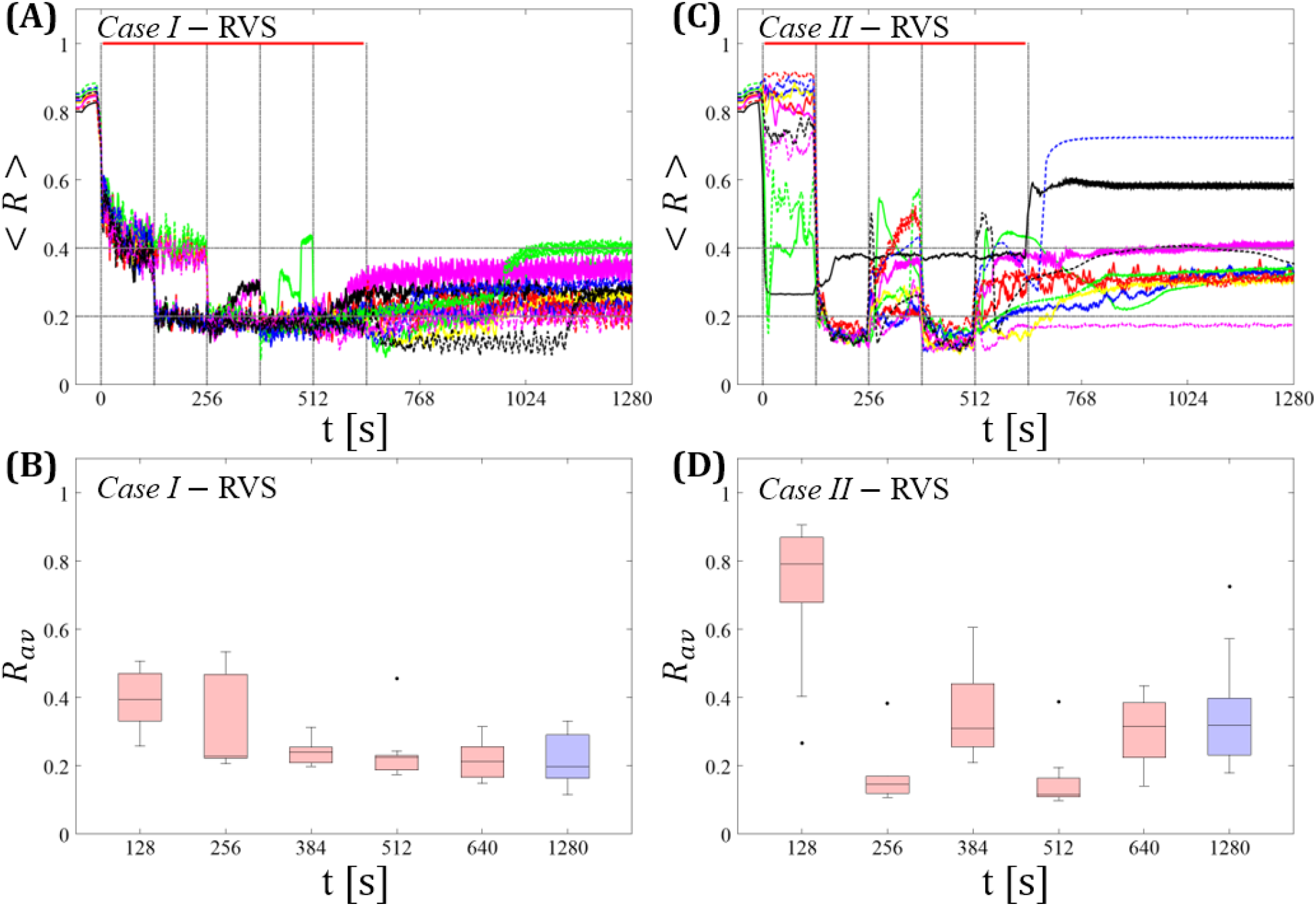
Long singleshot CR stimulation with demand-controlled variation of the stimulation frequency. **(A,C)** Time evolution of the order parameter < *R* > averaged over a sliding window during 5 consecutive RVS CR shots. If *R_av_*, the order parameter averaged over a window of length 100 · *T_s_* at the end of a shot, exceeds 0.4, the CR stimulation period of the subsequent RVS shot is decreased by *T_s_* → *T_s_* — 1 ms (see text). Stimulation parameters are unfavorable for anti-kindling in *Case I* **(A,B)** and *Case II* **(C,D)** (see text). **(B,D)** Boxplots for the time-averaged order parameter *R_av_* at the end of each shot (red color) and at the end of CR-off period (blue color), illustrate the overall outcome for all tested 11 networks. *Case I*: (*K, T_s_*) = (0.30,11). *Case II*: (*K, T_s_*) = (0.20,28). Same format as in **Figure 7**.

##### Long singleshot SVS CR stimulation with demand-controlled variation of the stimulation frequency

In *Case III* this protocol does not show any systematical improvement (**Figures 12A,12B**). In fact, this is partly due to the fact that in some cases the network gets trapped in an unfavorable parameter variation loop, bouncing between *T_s_* = 9 ms and *T_s_* = 10 ms. In *Case IV* (**Figure 12C,12D**) the global evolution is quite similar to the one found for *Protocol B* (**Figure 6C,6D**), i.e. pronounced desynchronization during a single shot, with a tendency to relapse back to the synchronized state while some of the networks remain desynchronized. However, the overall final outcome is rather poor as the corresponding boxplot (blue color) at the end of the CR-off period indicates.

**Figure 12.**
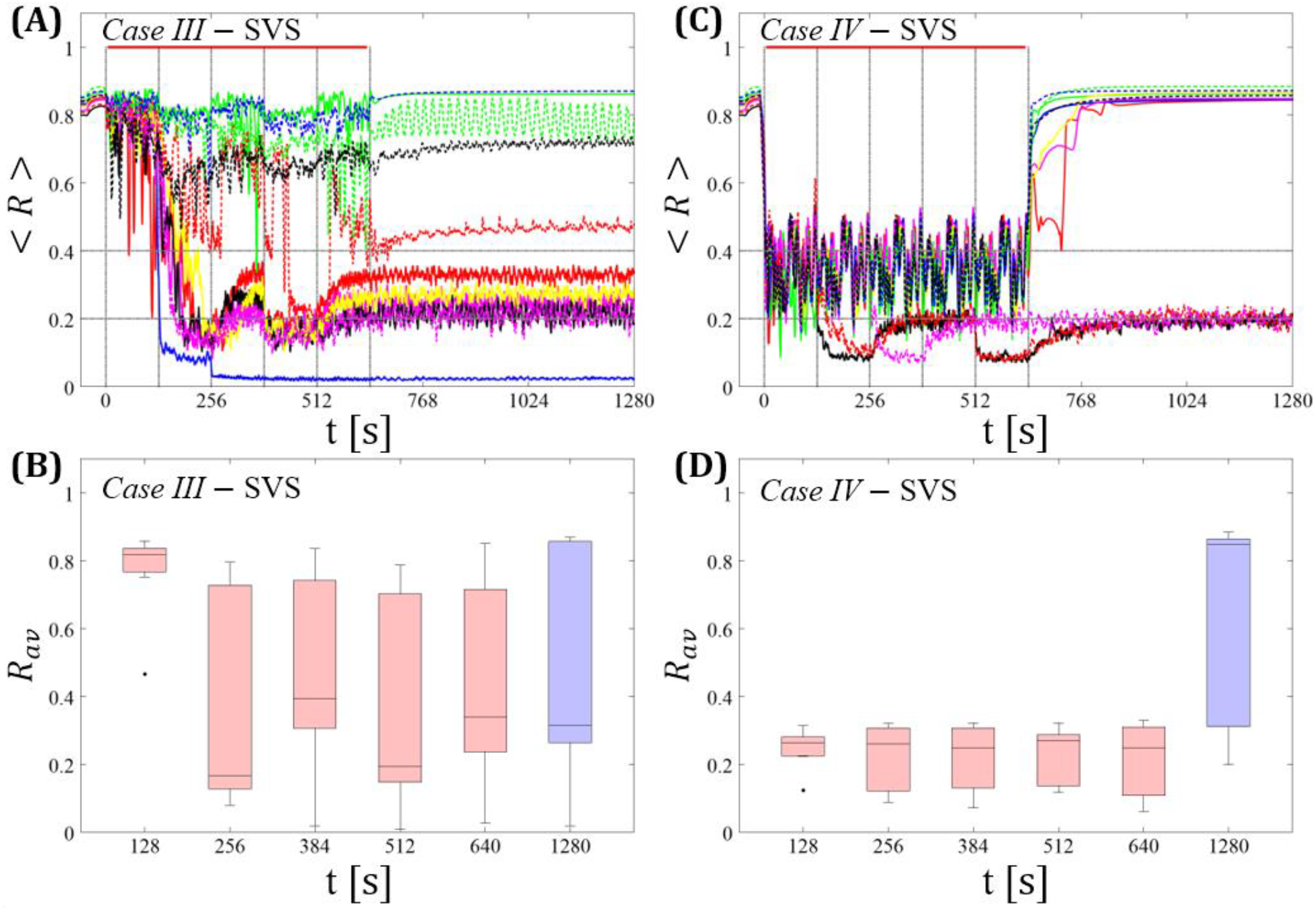
Long singleshot CR stimulation with demand-controlled variation of the stimulation frequency. **(A,C)** Time evolution of the order parameter < *R* > averaged over a sliding window during 5 consecutive RVS CR shots. If *R_av_*, the order parameter averaged over a window of length 100 · *T_s_* at the end of a shot, exceeds 0.4, the CR stimulation period of the subsequent RVS shot is decreased by *T_s_* → *T_s_* — 1 ms (see text). Stimulation parameters are unfavorable for anti-kindling in *Case III* **(A,B)** and *Case IV* **(C,D)** (see text). **(A,C)** The horizontal solid red lines indicate the CR shots, while the horizontal dashed grey lines serve as visual cues. **(B,D)** Boxplots for the time-averaged order parameter *R_av_* at the end of each shot (red color) and at the end of CR-off period (blue color), illustrate the overall outcome for all tested 11 networks. *Case* III: (*K, T_s_*) = (0.20,9). *Case IV*: (*K,T_s_*) = (0.20,14).

In summary, for the singleshot RVS CR as well as SVS CR stimulation *Protocol D* with regular variation of the CR stimulation duration *T_s_* (without pauses between two consecutive shots) did not lead to a reliable and systematic long-lasting desynchronization.

## Discussion

By comparing spaced CR stimulation with fixed stimulation parameters (*Protocol A*) and massed, continuous CR stimulation with equal integral duration (*Protocol B*) with a flexible spaced CR stimulation with demand-controlled variation of CR stimulation frequency and intensity (*Protocol C*), and with a flexible non-spaced CR stimulation with demand-controlled variation of CR stimulation frequency and intensity (*Protocol D*), we demonstrated that *Protocol C* enables to significantly improve the long-term desynchronization outcome of both RVS and SVS CR stimulation, even at comparatively short integral stimulation duration. Remarkably, spacing alone (*Protocol A*) is not sufficient to provide an efficient short-term dosage regimen (**Figures 2,3**). In fact, in particular cases fivefold longer stimulation duration might even be more efficient than five consecutive single CR shots with identical integral stimulation duration, at least for RVS CR stimulation (**Figures 2B** vs **4B**). The low performance of pure spacing (*Protocol A*) might be due to the low number of single CR shots, here five, as opposed to slightly larger numbers of CR shots, say eight, tested for the case of subcritical CR stimulation before (Popovych et al., 2015). However, more important might be the approx. fifty-fold longer stimulation and pause duration used for the spaced subcritical CR stimulation protocol (Popovych et al., 2015). The long spaced subcritical CR stimulation protocol might be beneficial for invasive application, such as DBS, and help reduce side-effects by substantially reducing stimulation current intake of the issue.

However, computationally we show that a spacing with rigid five-shot timing structure, but flexible, demand-controlled variation of stimulation frequency and intensity (*Protocol C*) provides a short-term dosage regimen that significantly improves the long-term desynchronization outcome of RVS and SVS CR stimulation (**Figures 6–9**). At the end of each pause between CR shots, the stimulus after-effect is assessed. If the desynchronization is considered to be insufficient, a mild variation of the CR stimulation frequency is performed to possibly provide a better fit between network and CR stimulation frequency, without actually adapting the stimulation frequency to frequency characteristics of the network stimulated. If desynchronization is considered to be moderate, the subsequent CR shot is delivered with parameters unchanged. If desynchronization is sufficient, CR stimulation is suspended during the subsequent shot. Intriguingly, in the vast majority of parameters and networks tested, this short-term dosage regimen induces a robust and reliable long-lasting desynchronization (**Figures 6–9**). This protocol might be a candidate especially for non-invasive, e.g. acoustic (Tass et al., 2012a) or vibrotactile (Syrkin-Nikolau et al., 2017;Tass, 2017), applications of CR stimulation to increase desynchronization efficacy, while keeping the stimulation duration at moderate levels.

Demand-controlled variation of CR stimulation frequency and intensity (*Protocol D*) alone (i.e. without inserting pauses) is not sufficient to significantly improve the outcome of RVS and SVS stimulation (**Figures 11,12**). Hence, introducing pauses significantly improves the effect of the demand-controlled variation of CR stimulation frequency and intensity.

In principle, stimulation parameters other than the CR stimulation frequency might be varied depending on the stimulation outcome. However, in this study we have chosen to vary the CR stimulation frequency, since the latter turned out to be a sensitive parameter, especially for SVS CR stimulation [see (Manos et al., 2017)]. In fact, the short-term dosage regimen with demand-controlled variation of stimulation parameters (*Protocol C*) might help to turn SVS CR stimulation in a method that causes a particularly strong anti-kindling in a robust and reliable manner.

*Protocol C* does not require a direct adaption of the CR stimulation frequency to measured quantities reflecting frequency characteristics of the network. We have chosen this design, since it might be an advantage not to rely on specific biomarker-type of information. For instance, in the case of PD a number of relevant studies were devoted to closed-loop DBS (Graupe et al., 2010;Rosin et al., 2011;Carron et al., 2013;Little et al., 2013;Priori et al., 2013;Yamamoto et al., 2013;Hosain et al., 2014;Rosa et al., 2015). A relevant issue in this context is the availability of a biomarker adequately reflecting the individual patient’s extent of symptoms (Beudel and Brown, 2016;Kühn and Volkmann, 2017). In fact, it is not clear whether low or high frequency beta band oscillations might be more appropriate as biomarker-type of feedback signal (Beudel and Brown, 2016). For several reasons, beta band oscillations might possibly not be an optimal feedback signal (Johnson et al., 2016;Kühn and Volkmann, 2017). Enhanced beta band oscillations are not consistently found in all PD patients (Kühn et al., 2008;Kühn and Volkmann, 2017). The clinical score of PD patients might more appropriately be reflected by the power ratio of two distinct bands of high frequency oscillations around 250 Hz and 350 Hz (Özkurt et al., 2011). Appropriate biomarkers might depend on the patient phenotype (Quinn et al., 2015): In tremor dominant (compared to akinetic rigid) PD patients resting state beta power may decrease during tremor epochs (Bronte-Stewart et al., 2009;Quinn et al., 2015). By a similar token, theta and beta oscillations interact with high-frequency oscillations under physiological (Yanagisawa et al., 2012) as well as pathological (Yang et al., 2014) conditions. Also, quantities assessing the interaction of brain oscillation, e.g. phase amplitude coupling (PAC) might be used as biomarker to represent the amount of symptoms (Beudel and Brown, 2016). Also, activity in the beta band might be relevant for compensatory purposes, as recently shown in a parkinsonian monkey study with sensorimotor rhythm neurofeedback (Philippens et al., 2017).

It might be another potential advantage for clinical applications that the three-stage control of the proposed short-term dosage regimen (*Protocol C*) could possibly be approximated by scores reflecting the patient’s state or the amount symptoms. A simple three-stage rating of the patient’s state (bad, medium, and good) might replace the feedback signal-based stages (i), (ii), and (iii). Assessments of the patient’s state might be performed in a pause after a CR shot. Depending on the rating, the CR stimulation frequency or intensity of the subsequent CR shot may be varied. In particular, for non-invasive application of CR stimulation a non-invasive assessment of the stimulation effect might straightforwardly be realized.

In realistic biological systems intrinsic (model) parameters typically vary over time. These variations may be of complex dynamical nature [see e.g. (Timmer et al., 2000;Yulmetyev et al., 2006)]. To obtain some indication as to whether *Protocol C* is robust against low-amplitude intrinsic variations of the neuronal firing rates, we added a low-amplitude term *I_var_* = *A* · *sin*(2*π* · *f · t*) to the right-hand side of Equation 1a. In the stimulation-free case, *I_var_* causes variations of the neurons’ firing rates in the order of ±3% and no qualitative changes of the network dynamics (data not shown). For different frequencies *f* this type of variation does not significantly affect the long-term desynchronization outcome of *Protocol C* (*f* = 0.004 Hz, 4 Hz and 20 Hz in **Supplementary Figure 1**). By the same token, the neuronal firing rates are not significantly altered by the additional periodic force (data not shown).

Note, this is not intended to be a comprehensive study of the impact of periodic forcing of arbitrary frequency on the spontaneous or stimulation-induced dynamics of the model network under consideration. Rather, the slow oscillatory forcing is meant to model slow physiological modulatory processes in an illustrative manner. In the extreme case of *f* = 0.004 Hz the slow oscillatory modulation acts on the same time scale as a cycle comprising shot and pause and, hence smoothly emulates the step-wise modulation of the CR stimulation frequency in *Protocol C*.

Conversely, intrinsic variations of sufficient size might naturally mimic variations of the relationship between CR stimulation frequency and intrinsic neuronal firing rates as introduced on purpose in *Protocol C*. Accordingly, already the purely spaced stimulation without demand-controlled variability (*Protocol A*) might display some variability of the relationships between intrinsic firing rates and CR stimulation frequency simply due to the intrinsic variability. However, at least with the frequencies 0.004 Hz, 4 Hz and 20 Hz in the low-amplitude term *I_var_* = *A* · *sin*(2*π* · *f · t*) added to the right-hand side of Equation 1a, we were not able to observe any substantial improvement of the desynchronizing outcome of *Protocol A* (**Supplementary Figure 2**). However, more physiological patterns of firing rate modulations might have a more significant impact on the stimulation outcome of *Protocol A*. In future studies typical variations of the signals relevant to a particular pre-clinical or clinical application might be taken into account to further improve desynchronizing short-term dosage regimen. The additional periodic forcing considered here was meant to illustrate the stability of the suggested control approach. However, future studies could also provide a detailed analysis of the interplay of one or more periodic inputs and noise, thereby focusing on stochastic resonance and related phenomena [e.g. (Pikovsky and Kurths, 1997;Gammaitoni et al., 1998;Manjarrez et al., 2002;Torres et al., 2011;Bordet et al., 2015;Yu et al., 2016;Guo et al., 2017;Uzuntarla et al., 2017) and references therein].

The short-term dosage regimen proposed here provides a closed-loop CR stimulation concept that enables to significantly increase the robustness and reliability of the stimulation outcome. Our results motivate to further improve the CR approach by closed loop or feedback-based dosage regimen. Compared to the computationally developed initial concept of demand-controlled CR-induced desynchronization of networks with fixed coupling constants (Tass, 2003a;Tass, 2003b), the focus will now be on a feedback-adjusted modulation of synaptic patterns to induce long-lasting therapeutic effects. Currently, clinical proof of concept (phase IIa) is available for deep brain CR stimulation for the therapy of Parkinson’s disease (Adamchic et al., 2014) and acoustic CR stimulation for the treatment of chronic subjective tinnitus (Tass et al., 2012a). In addition, promising first in human (phase I) data are available for vibrotactile CR stimulation for the treatment of Parkinson’s disease showing pronounced and highly significant sustained therapeutic effects (Syrkin-Nikolau et al., 2017). For the clinical development of these treatments it is mandatory to perform dose-finding studies (phase IIb) to reveal optimal stimulation parameters and dosage regimens, for comparison see (Friedman et al., 2010). The latter are required to get properly prepared for large efficacy (phase III) trials (Friedman et al., 2010). Since CR stimulation modulates complex neuronal dynamics, dose-finding studies are sophisticated, since stimulation parameters as well as dosage patterns have to be chosen appropriately. Selecting appropriate stimulation parameters and dosage regimens by trial and error may neither be effective nor affordable, since it would require a huge number of patients. In contrast, our manuscript illustrates the important role of computational medicine in generating hypotheses for dose-finding studies. Specifically, we show that spacing (i.e. adding pauses in between stimulation epochs) as well as moderate and unspecific parameter variations adapted in the course of the therapy are not sufficient to overcome limitations of CR stimulation. Intriguingly, the combination of both, spacing plus adaptive moderate parameter variation increases the robustness of the stimulation outcome in a significant manner. This computational prediction can immediately be tested in dose-finding studies and, hence, help to optimize the CR therapy, shorten the development time and reduce related costs.

## Author Contributions

Conceived and designed the experiments: TM MZ PAT. Performed the experiments: TM. Analyzed the data: TM MZ PAT. Contributed reagents/materials/analysis tools: TM MZ. Wrote the paper: TM MZ PAT.

## Supplementary Material

**Supplementary Figure 1.**
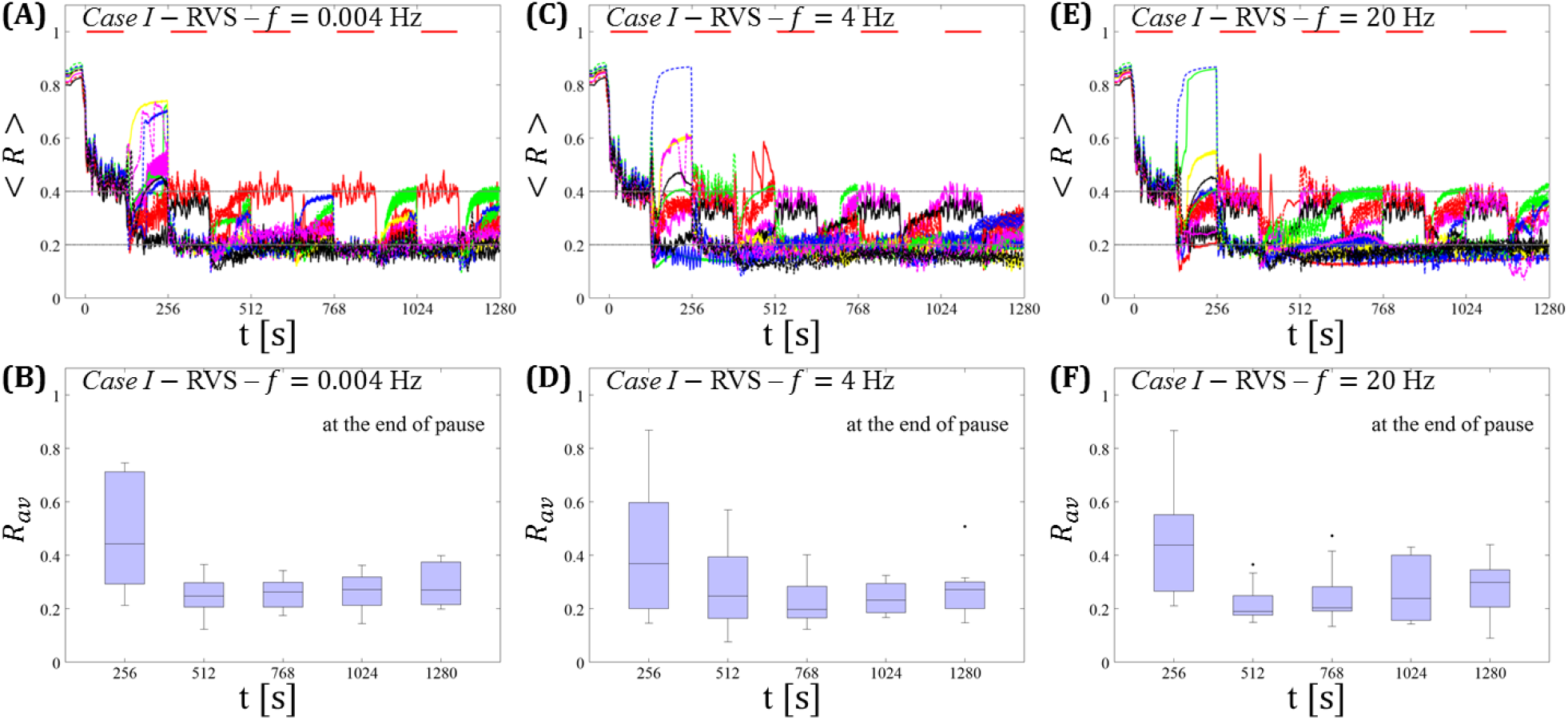
Protocol C in the presence of intrinsic variations of the firing rates caused by a modulatory low-amplitude current input *I_var_* = *A* · *sin*(2*π · f · t*), with *A* = 1, *f* = 0. 004 Hz (A, B), *f* = 4 Hz (C, D) and *f* = 20 Hz (E, F). Spaced multishot RVS CR stimulation with demand-controlled random variation of the stimulation period *T_s_* and with demand-controlled variation of the intensity. The low-amplitude variation *I_var_* is active during the entire simulations, respectively. Its low amplitude ***A*** = 1 ensures that the dynamics of the network is not drastically affected. **(A, C, E)** Time evolution of the order parameter < *R* > averaged over a sliding window during 5 consecutive RVS CR shots respectively. If *R_av_* at the end of a pause exceeds 0.4, the CR stimulation period of the subsequent SVS shot is decreased by *T_s_* → *T_s_* − 1 ms (see text). **(B, D, F)** Boxplots for the time-averaged order parameter *R_av_* at the end of each pause, illustrate the overall outcome for all tested 11 networks respectively. The horizontal solid red lines indicate the CR shots, while the horizontal dashed grey lines highlight the two control thresholds (see text). *Case I* stimulation parameters are unfavourable for anti-kindling: (*K, T_s_*) = (0.30,11) (see text). Format as in **Figure 9**.

**Supplementary Figure 2.**
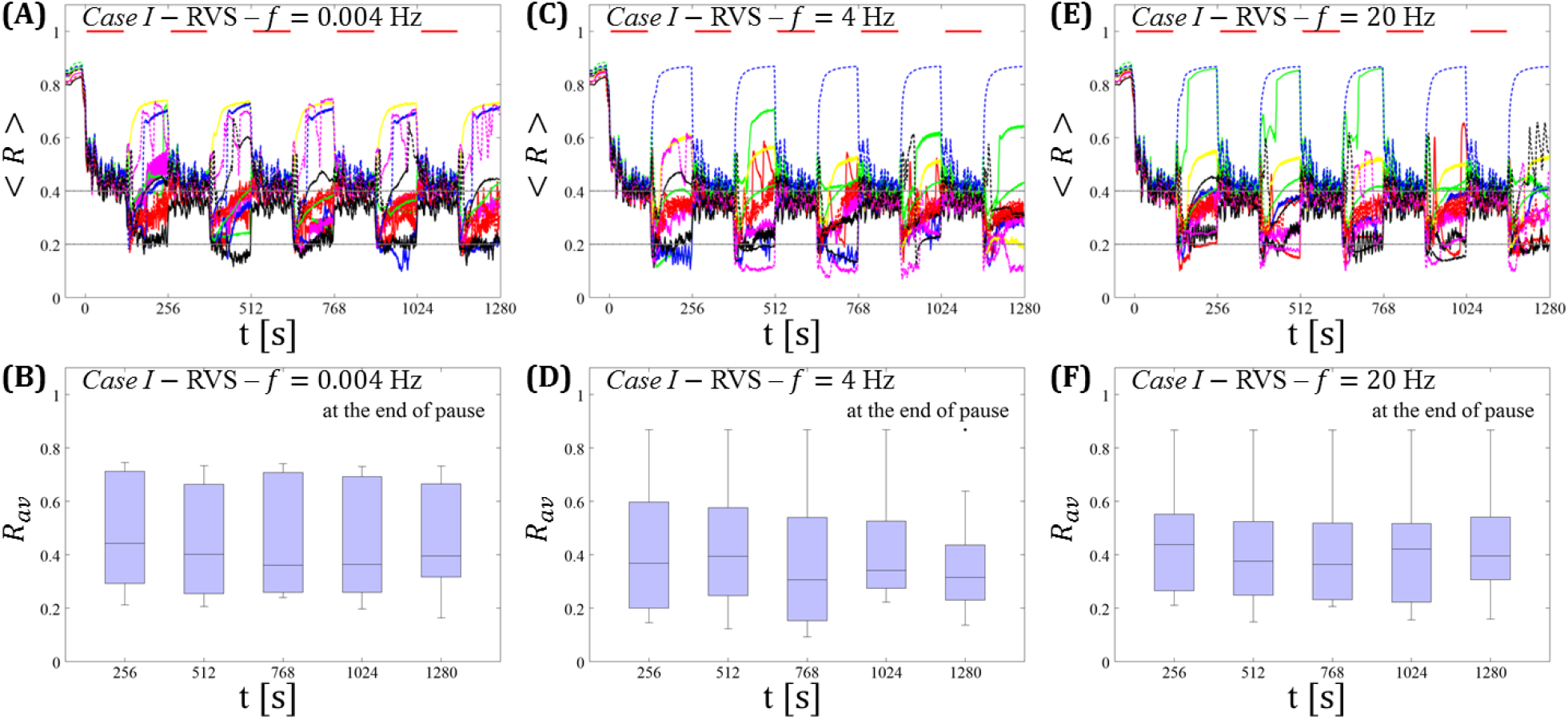
Protocol A in the presence of intrinsic variations of the firing rates caused by a modulatory low-amplitude current input *I_var_* = *A* · *sin*(2*π · f · t*), with *A* = 1, *f* = 0. 004 Hz **(A, B)**, *f* = 4 Hz **(C, D)** and *f* = 20 Hz **(E, F)**. Spaced multishot RVS CR stimulation with fixed stimulation period *T_s_*. Same series of simulations and analysis as in **Supplementary Figure 1**. **(A, C, E)** Time evolution of the order parameter < *R* > averaged over a sliding window during 5 consecutive RVS CR shots respectively. **(B, D, F)** Boxplots for the time-averaged order parameter *R_av_* at the end of each pause, illustrate the overall outcome for all tested 11 networks respectively. Spacing is symmetrical, i.e. CR shots and consecutive pauses are of the same duration. *Case I* stimulation parameters are unfavourable for anti-kindling: (*K, T_s_*) = (0.30,11) (see text). Format as in **Figure 9**.

